# Massive haplotypes underlie ecotypic differentiation in sunflowers

**DOI:** 10.1101/790279

**Authors:** Marco Todesco, Gregory L. Owens, Natalia Bercovich, Jean-Sébastien Légaré, Shaghayegh Soudi, Dylan O. Burge, Kaichi Huang, Katherine L. Ostevik, Emily B. M. Drummond, Ivana Imerovski, Kathryn Lande, Mariana A. Pascual, Winnie Cheung, S. Evan Staton, Stéphane Muños, Rasmus Nielsen, Lisa A. Donovan, John M. Burke, Sam Yeaman, Loren H. Rieseberg

**Author notes:** These authors made similar contributions to this work (Marco Todesco, Gregory L. Owens, Natalia Bercovich). Corresponding authors: Correspondence and requests for material should be addressed to (L.H.R.), (N.B.).

## Abstract

Species often include multiple ecotypes that are adapted to different environments. But how do ecotypes arise, and how are their distinctive combinations of adaptive alleles maintained despite hybridization with non-adapted populations? Re-sequencing of 1506 wild sunflowers from three species identified 37 large (1-100 Mbp), non-recombining haplotype blocks associated with numerous ecologically relevant traits, and soil and climate characteristics. Limited recombination in these regions keeps adaptive alleles together, and we find that they differentiate several sunflower ecotypes; for example, they control a 77 day difference in flowering between ecotypes of silverleaf sunflower (likely through deletion of a *FLOWERING LOCUS T* homolog), and are associated with seed size, flowering time and soil fertility in dune-adapted sunflowers. These haplotypes are highly divergent, associated with polymorphic structural variants, and often appear to represent introgressions from other, possibly extinct, congeners. This work highlights a pervasive role of structural variation in maintaining complex ecotypic adaptation.

---

Local adaptation is common in species that experience different environments across their range. This can result in the formation of ecotypes, ecological races with distinct morphological and/or physiological characteristics that provide an environment-specific fitness advantage. Despite the prevalence of ecotypic differentiation, much remains to be understood about its genetic basis and the evolutionary mechanisms leading to its establishment and maintenance. In particular, a longstanding evolutionary question, dating back to criticisms of Darwin’s theories by his contemporaries^1^, concerns how such ecological divergence can occur in the presence of hybridization with non-adapted populations^2^. Local adaptation typically requires alleles at multiple loci contributing to increased fitness in the same environment; however, different ecotypes are often geographically close and interfertile, and gene flow between them should break up adaptive allelic combinations by promoting recombination with non-adaptive alleles^3^.

To better understand the genetic basis of local adaptation and ecotypic differentiation, we conducted an in-depth study of genetic, phenotypic, and environmental variation in three annual sunflower species, which include multiple reproductively compatible ecotypes. Two of these species have broad, overlapping distributions across North America; *Helianthus annuus*, the common sunflower, is the closest wild relative of cultivated sunflower, which was domesticated from it around 4,000 years ago in East-Central North America^4^. Populations of *H. annuus* are generally found on mesic soils, but can grow in a variety of disturbed or extreme habitats, including semi-desertic or frequently flooded areas, as well as salt marshes. An especially well-characterized ecotype (formally *H. annuus* subsp. *texanus*), is adapted to the higher temperatures and herbivore pressures in Texas, USA^5^. *Helianthus petiolaris*, the prairie sunflower, prefers sandier soils, and ecotypes of this species are adapted to sand sheets and dunes^6^. The third species, *Helianthus argophyllus*, the silverleaf sunflower, is found exclusively in southern Texas and includes both an early flowering coastal island ecotype and a late flowering inland ecotype^7^.

## Population structure of wild sunflowers

We grew in a common garden experiment ten plants from each of 151 populations of these three sunflower species (*H. annuus* = 71 populations; *H. petiolaris* = 50 populations; *H. argophyllus* = 30 populations), selected from across their native range (Fig. 1a), and for which we collected corresponding soil samples. We generated extensive records of developmental and morphological traits throughout the growth of the plants, and re-sequenced the genome of 1401 of them. An additional 105 *H. annuus* individuals were re-sequenced to fill gaps in geographic coverage, as well as twelve annual and perennial sunflowers to be used as outgroups (Supplementary Table 1). Sunflower genomes are relatively large (*H. annuus* = 3.5 Gbp; *H. petiolaris* = 3.3 Gbp; *H. argophyllus* = 4.3 Gbp^8^) and comprised of >75% retrotransposon sequences^9^. We used enzymatic depletion^10^ to reduce the proportion of repetitive sequences, resulting in an average 6.34-fold coverage of gene space (median = 6.03; Supplementary Table 1). Sequencing reads were aligned to the reference genome of cultivated sunflower^9, 11, 12^ (Extended Data Fig. 1), resulting in sets of >4M high-quality single nucleotide polymorphisms (SNPs) for each species.

**Fig. 1:**
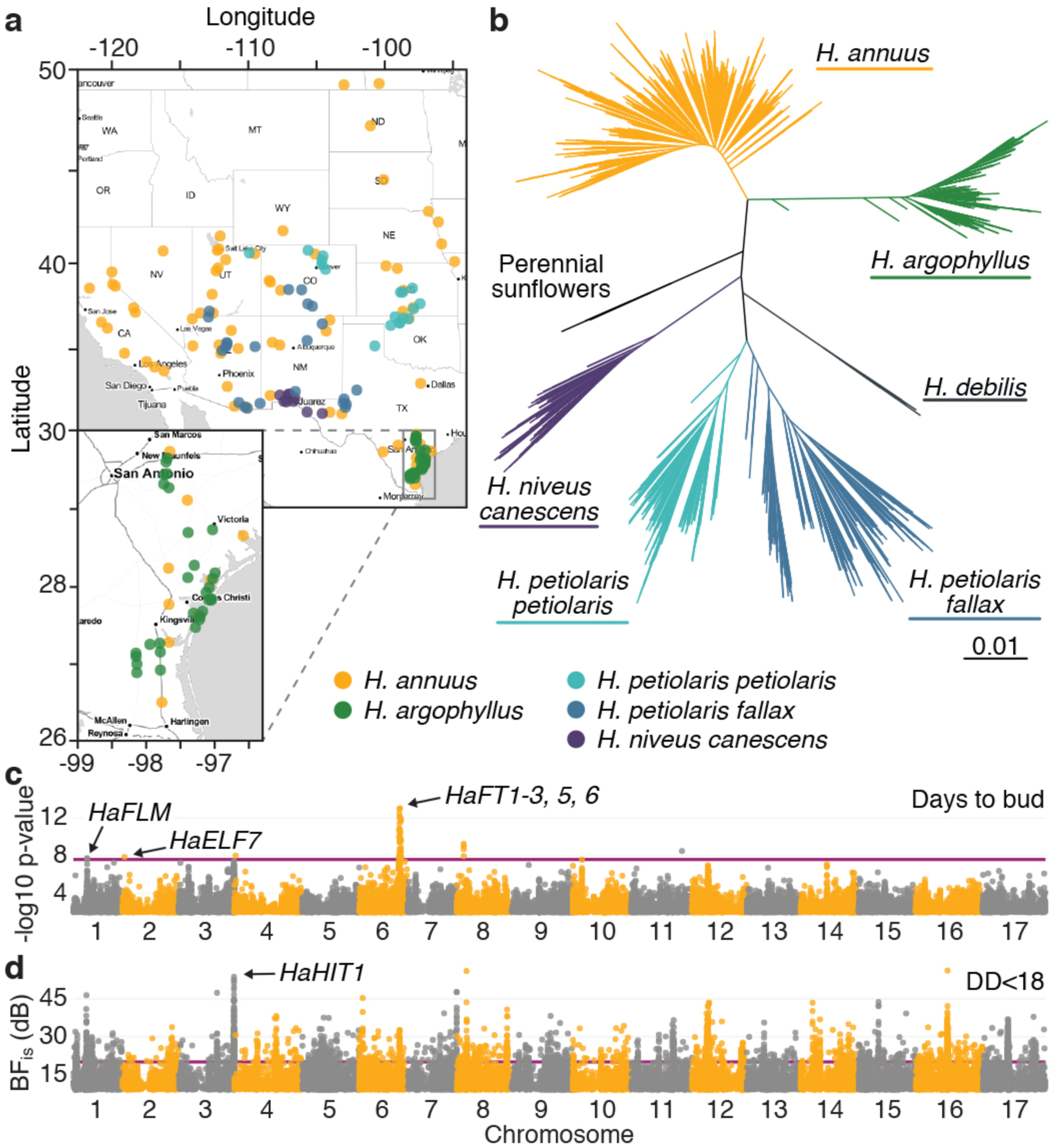
Population structure and association analyses of wild sunflowers. **A**, Map of wild sunflower populations surveyed in this study. **b**, Maximum-likelihood tree for the samples included in this study. **C**, Flowering time GWA for *H. annuus*. The purple line represents 5% significance after Bonferroni correction. Only positions with -log10 p-value > 2 are plotted. **D**, Degree-days below 18 °C (DD<18) GEA for *H. annuus*. The purple line represents Bayes Factor (BF_is_) = 20 deciban (dB). Only positions with BF_is_ > 9 dB are plotted.

A phylogeny based on these and previously re-sequenced sunflower samples agrees with earlier studies^13, 14^: *H. annuus* and *H. argophyllus* are sister species, whereas *H. petiolaris* is placed in a separate clade. We found three separate lineages within our *H. petiolaris* collection, corresponding to the subspecies *fallax, petiolaris*, and *canescens*. However, subsp*. canescens* falls within *H. niveus*, supporting an earlier classification^15^; due to its smaller sample size (86 individuals), we have omitted the *H. niveus canescens* clade from further analyses. Lastly, dune-adapted ecotypes of *H. petiolaris* from Colorado and Texas fall within *H. petiolaris fallax* (despite the Texas populations being formally designed as *H. neglectus*^16^), and are therefore analyzed as part of that clade (Fig. 1b).

## Large haplotypes linked to adaptive traits

The large effective population size and outcrossing mating system of wild sunflowers^17^ represents a major advantage for genome-wide association (GWA) studies, since the rapid decay of linkage disequilibrium (LD) permits mapping of phenotype-genotype associations to narrow genomic regions if (as in this case) sufficient marker densities are available. GWA analyses of 87 traits identified numerous, strong links between phenotypic variation and regions of the sunflower genome (Supplementary Table 2). We observed, for example, extensive variation in flowering time for all three species (Extended Data Fig. 2a), consistent with its fundamental role in plant (and sunflower) adaptation^18, 19^. For *H. annuus* in particular, significant associations were found with the sunflower homologs of known flowering time regulators, including *FLOWERING LOCUS T*^20^ (*FT*), *FLOWERING LOCUS M*^21^ (*FLM*), and *EARLY FLOWERING 7*^22^ (*ELF7*; Fig. 1c). We also identified several regions of the genome strongly associated with environmental and soil variables in genotype-environment association (GEA) analyses, suggesting a role in adaptation to particular habitats (Supplementary Table 2). For example, several temperature-related variables showed strong associations with the sunflower homolog of *HEAT-INTOLERANT 1* (*HIT1*), which mediates resistance to heat stress by regulating plasma membrane thermo-tolerance in *Arabidopsis thaliana*^23^ (Fig. 1d).

In several cases, we noticed GWA and GEA signals spanning very large regions of the genome for traits that are known to be important for local adaptation, and to differentiate ecotypes in sunflower. One of the most striking examples of these GWA plateaus occurred between coastal island and inland populations of *H. argophyllus*. While inland populations flower late in summer, and can grow to be extremely large (up to >4 meters), smaller early flowering individuals occur at high frequency on the barrier islands of the Gulf of Mexico (Fig. 2a,b). Selection experiments indicate that late flowering in the interior is favoured^7^, presumably to avoid flowering during the extremely hot and dry summer, whereas early flowering appears to be advantageous under less harsh conditions on the barrier islands. Flowering time GWA analyses in *H. argophyllus* identified a single, highly significant association spanning ∼30 Mbp on chromosome 6 (Fig. 2c,d), also associated with leaf nitrogen and carbon content (Extended Data Fig. 2b). Principal component analysis (PCA) of this region suggested the presence of two main haplotypes, with intermediate individuals being heterozygotes (Fig. 2e). We extracted haplotype-informative sites and visualized ancestry across the region, revealing very limited recombination. A 10 Mbp region (130-140 Mbp) is perfectly correlated with flowering time phenotypes and explains 88.2% of the variance in days to bud (Fig. 2f). The early haplotype acts dominantly, with plants carrying at least one copy of it flowering on average 77 days earlier than late-flowering individuals (Fig. 2g). This region contains five of the six sunflower homologs of the flowering time regulator *FT* (*HaFT1-3*, *HaFT5*, *HaFT6*; Fig. 2f). Surprisingly, the GWA signal drops sharply around the *HaFT1* locus (Fig. 2d), which is known to play a role in differences in photoperiodic responses between wild and cultivated sunflower^24^. Analysis of an unfiltered SNP dataset revealed that this pattern is due to the almost complete absence of reads mapping to the region in plants carrying the late-flowering haplotype (only SNPs with data for at least 90% of the individuals were used for GWAs). This is consistent with the presence of one or more deletions, including the *HaFT1* locus, in late flowering *H. argophyllus* (Fig. 2h; Extended Data Fig. 2c). Accordingly, the *HaFT1* sequence cannot be amplified from genomic DNA from late-flowering plants, and no *HaFT1* expression is detected in those plants (Fig. 2i,j; Extended Data Fig. 2d). Early flowering plants carry instead at least one functional copy of *HaFT1*, which complements the otherwise late-flowering *A. thaliana ft*-10 mutant (Fig. 2k,l). To explore the origins of these haplotypes, we constructed a phylogeny of the non-recombined 10 Mbp region in chromosome 6 (Fig. 2m). We found that the two haplotypes are highly divergent, and that the early haplotype was introgressed from *H. annuus* (D-stat = 0.844 ∓ 0.006, p < 10^-20^, two-sided; see also Fig. 2g). While it is not possible to exclude a role of the other *FT* homologs (Extended Data Fig. 2e-g) or other genes in the region, these results strongly suggest that introgression of a functional *HaFT1* copy from *H. annuus* played a major role in the establishment of early-flowering *H. argophyllus*.

**Fig. 2:**
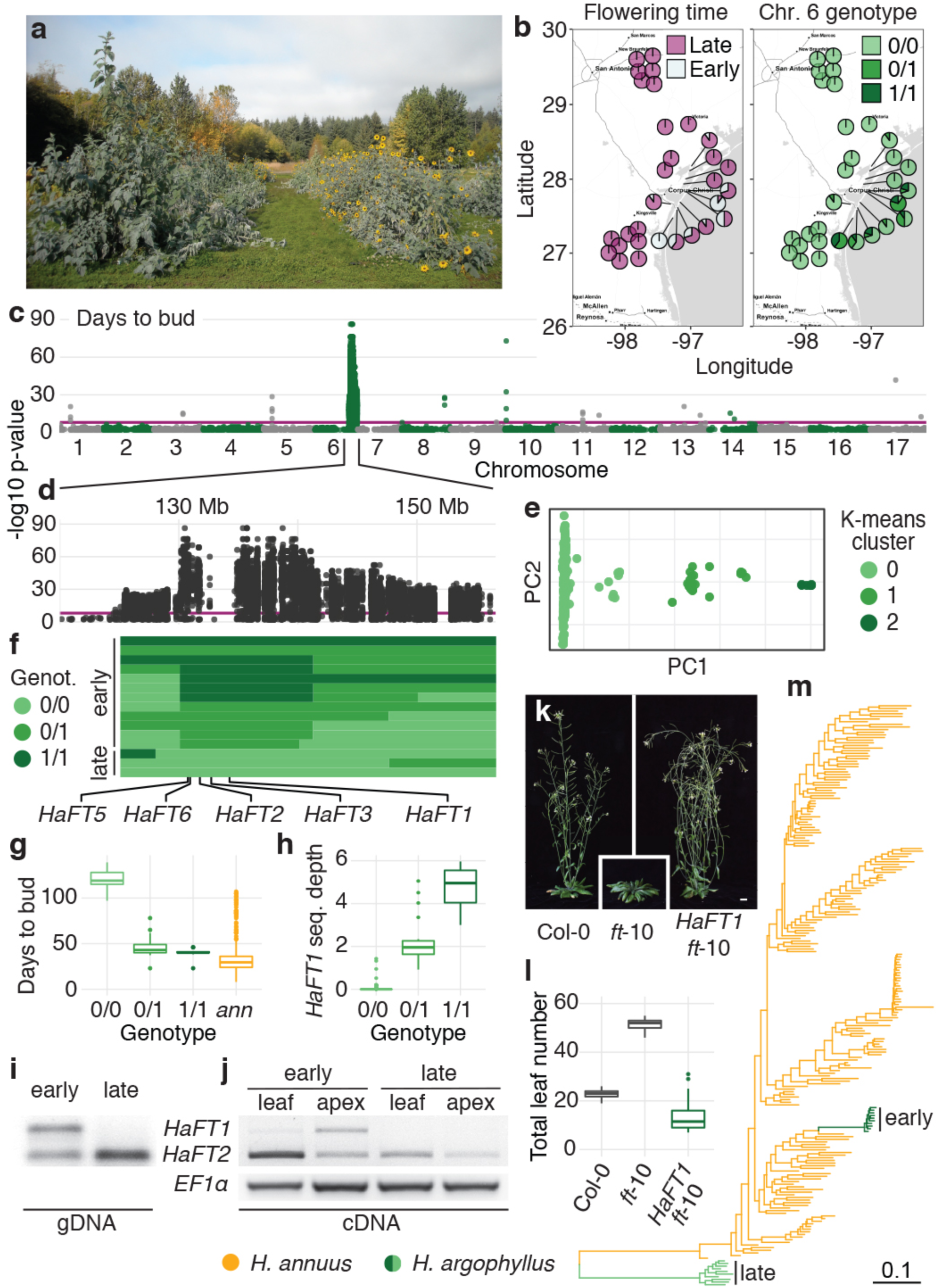
Early flowering in coastal *H. argophyllus* ecotypes is due to introgression from *H. annuus* of a large region containing a functional copy of *HaFT1*. **a**, Individuals from inland (left) and coastal island (right) populations of *H. argophyllus*. **b**, Geographic distribution of late- and early-flowering ecotypes (left), and of haplotypes of the causal 10 Mbp region on chromosome 6 (right), across populations of *H. argophyllus*. **c**, Flowering time GWA in *H. argophyllus* and **d**, zoomed view of the bottom of chromosome 6. The purple line represents 5% significance after Bonferroni correction. Only positions with - log10 p-value > 2 are plotted. **e**, PCA of the last 30 Mbp of chromosome 6. Three clusters are defined by PC1. **f**, Schematic representation of recombinant haplotypes, divided based on whether they are associated with early or late flowering. Only unique haplotypes are presented. Chromosome positions match panel d**. g**, Flowering time for individuals with different genotypes at the ∼130-140 Mbp region of chromosome 6. *ann* = *H. annuus*. **h**, Sequencing depth of SNPs in the *HaFT1* gene. Sequencing depth was comparable across haplotypes for the other four *HaFT* genes on chromosome 6. **i**, PCR on genomic DNA from early- and late-flowering *H. argophyllus* plants. **j**, Expression analysis in mature leaves or shoot apices of the plants examined in i, grown in long days conditions (14 hours light, 10 hours dark), at six weeks after planting. Cleaved-amplified polymorphic sequence (CAPS) markers were used to distinguish *HaFT1* from *HaFT2*. Experiments in panels i,j were repeated on three independent pairs of individuals, with similar results. **k**, Six-week-old *A. thaliana* plants grown in long day conditions. *ft*-10 plants carry a loss-of-function allele of the *AtFT* gene and flower extremely late in these conditions. Size bar = 1 cm. **l**, Flowering time (measured as total leaf number at flowering) for primary transformants expressing the *H. argophyllus HaFT1* gene in *ft*-10 background (n ≥ 25). Box plots show the median, box edges represent the 25^th^ and 75^th^ percentiles, whiskers represent the maximum/minimum data points within 1.5x interquartile range outside box edges. Differences between genotypes are significant for p < 10^-8^ (one-way ANOVA with post-hoc Tukey HSD test, F = 596, df = 2). **m**, Maximum likelihood phylogeny of the 130-140 Mbp region on chromosome 6 in *H. argophyllus* and *H. annuus*.

We found another clear example of GWA and GEA plateaus underlying ecotypic differentiation in *H. petiolaris*, which has repeatedly adapted to sand dunes in Texas and Colorado, USA^6^. Dune populations exhibit starkly distinctive phenotypes compared to populations growing just off the same dunes (Fig. 3a-d), the most striking of which are seed size and length (Fig. 3b,c); large seeds confer a strong fitness advantage on sand dunes^6^, possibly by providing seedlings with enough resources to emerge after being buried by sand. Dunes also are low in resources, and dune sunflowers use soil nutrients more efficiently that their non-dune counterparts^25^. GWA analyses for seed size and flowering time, and GEA analyses of soil characteristics including cation exchange capacity (CEC, a measure of soil fertility) in *H. petiolaris fallax* identified three multi-Mbp regions on chromosomes 9, 11 and 14 (Fig. 3d,e; Extended Data Fig. 3a,b). The plateaus on chromosomes 11 and 14 co-localize with known QTL for seed size differentiating dune and non-dune populations^26^, and weaker associations with flowering time are observed in *H. petiolaris petiolaris* for the chromosome 9 and 11 regions (Extended Data Fig. 3c). All three regions are highly differentiated between dune and non-dune populations from Texas, and two of the three regions differentiate dune and non-dune populations in Colorado^27^ (Fig. 3f), suggesting a fundamental role in maintaining the dune ecotype.

**Fig. 3:**
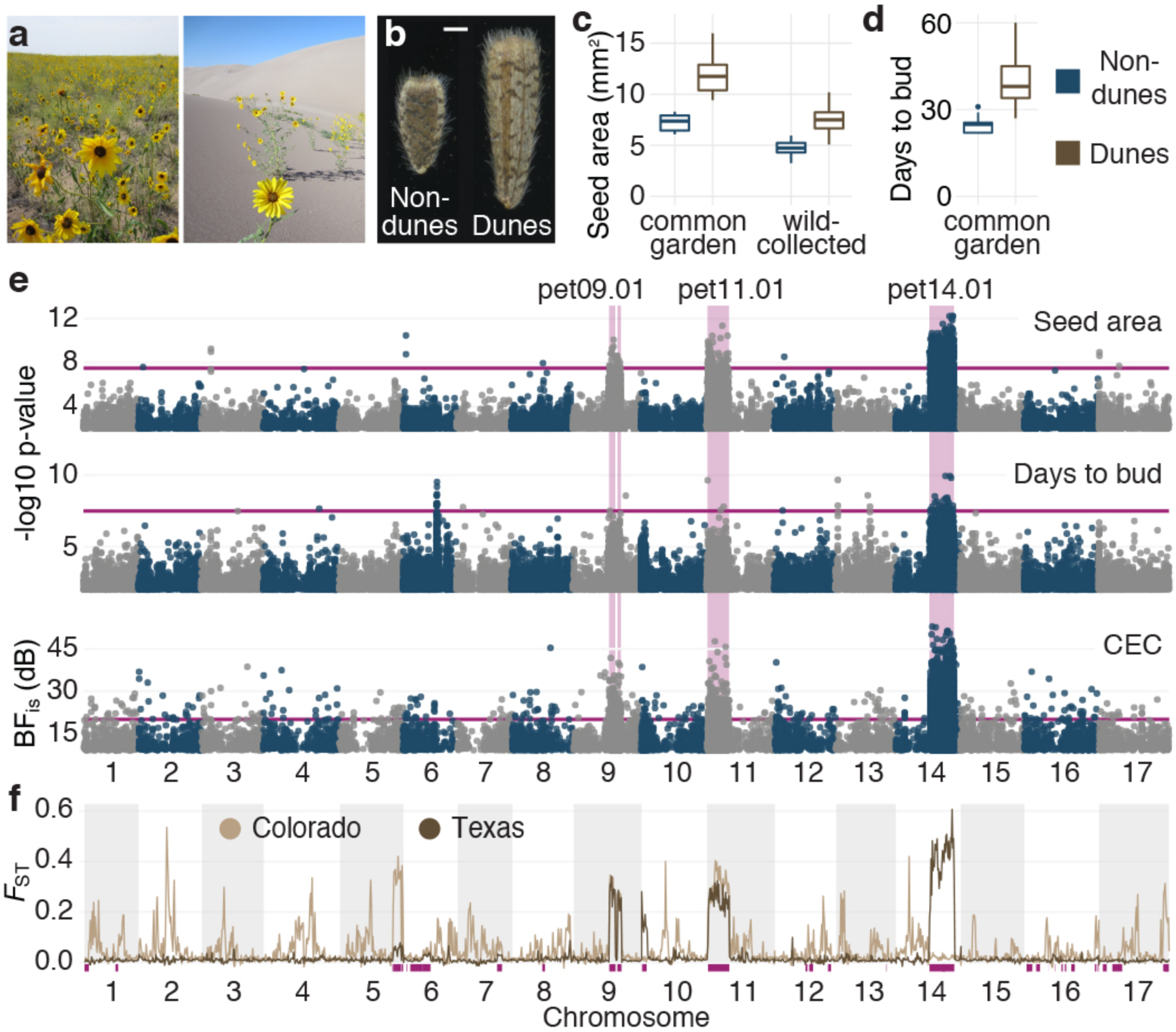
Large non-recombining haplotypes control dune adaptation in *H. petiolaris fallax*. **A**, *H. petiolaris fallax* populations growing on the sand sheet (left) or on sand dunes (right). **B**, Representative seeds from dune-adapted and non-dune-adapted plants grown in a common garden. Size bar = 1 mm. **c**, Seed size (area), averaged for eight seeds/plant. Seeds were collected in the wild or from plants grown in a common garden. While wild-collected seeds are generally smaller, seeds from dune-adapted plants are ∼60% larger in both comparisons (n ≥ 10; p < 10^-6^, Mann-Whitney U-test). **d**, Flowering time in a common garden (n ≥ 15; p < 10^-7^, Mann-Whitney U-test). Box plots show the median, box edges represent the 25^th^ and 75^th^ percentiles, whiskers represent the maximum/minimum data points within 1.5x interquartile range outside box edges. **e**, Seed size and flowering time GWAs and CEC (soil fertility) GEA for *H. petiolaris fallax*. Purple lines represent 5% significance after Bonferroni correction in GWA plots, and BF_is_= 20 dB in the GEA plot. Only positions with BF_is_ > 9 dB or -log10 p-value > 2 are plotted. Three haploblock predictions corresponding to three significant plateaus are highlighted in purple. **f**, F_ST_ values in 2 Mbp non-overlapping sliding windows for comparisons between dune- and non-dune-adapted populations of *H. petiolaris fallax* in Colorado and Texas. Purple bars represent predicted haploblocks. Some predicted haploblocks are in multiple pieces due to rearrangements in *H. petiolaris* relative to the *H. annuus* reference genome (Extended Data Fig. 6c).

## Highly divergent haploblocks are common

The identification of these GWA/GEA plateaus suggests a broader role of large, non-recombining haplotype blocks (henceforth “haploblocks”) in adaptation. To test this hypothesis, we used a local PCA approach to identify other large genomic regions with distinct population structure^28^ (Fig. 4a). Across the three species, we found 37 such regions, ranging from 1 to 100 Mbp and representing 4-16% of the total genome (Fig. 4b; Extended Data Table 1). They are characterized by high LD, and PCAs in these regions separated individuals into three clusters, with the middle cluster having higher heterozygosity. This is consistent with the two extreme clusters representing individuals homozygous for two distinct haplotypes, and the middle cluster representing heterozygotes. No or very little recombination is observed between haplotypes, but generally no reduction in recombination is found within haplotypes (Fig. 4c-e; Extended Data Fig. 4,5).

**Fig. 4:**
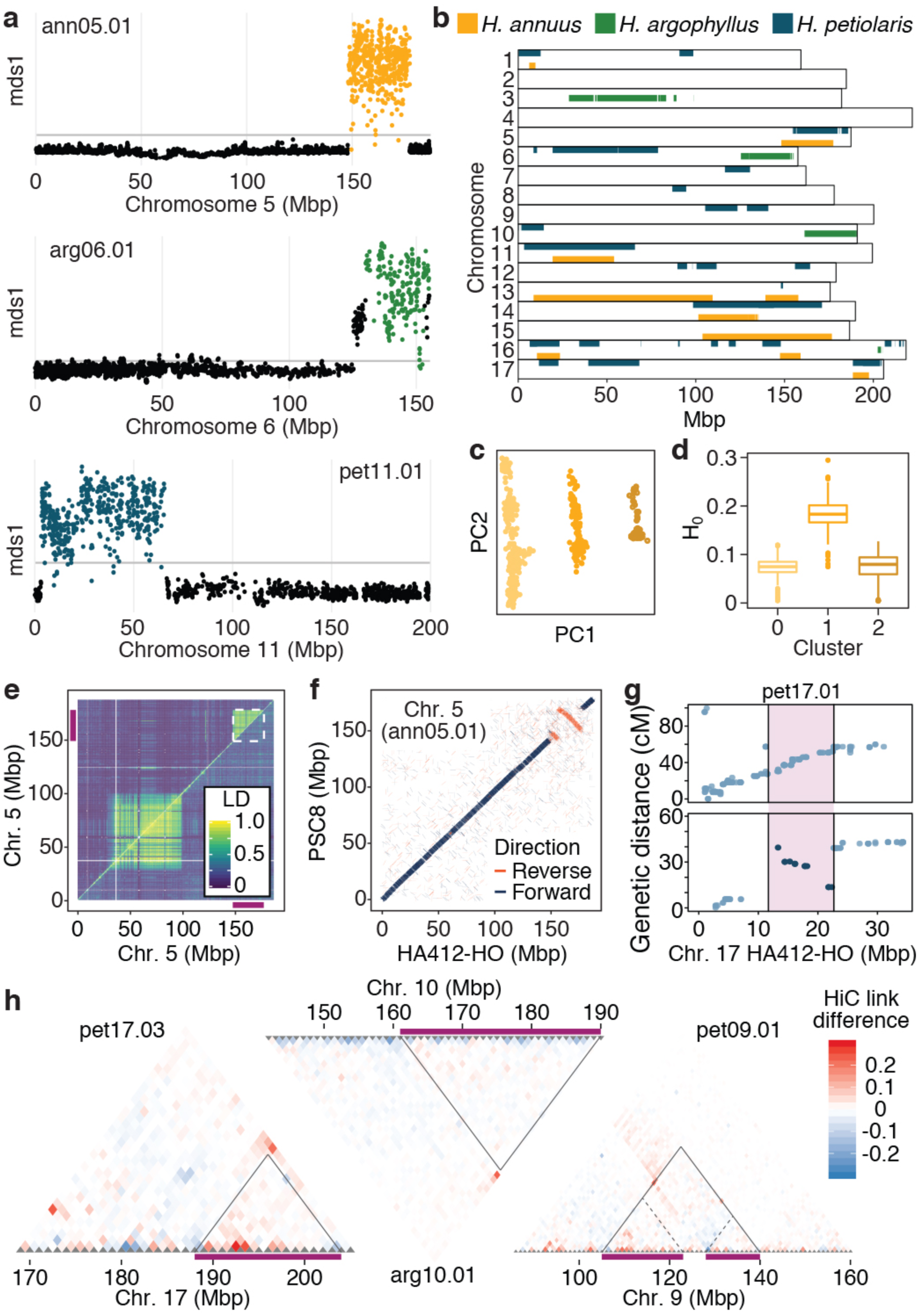
Large haploblocks are pervasive in wild sunflowers and are associated with SVs. **a**, Example local PCA outputs for putative haploblocks segregating in the three wild sunflower species. Each dot represents a 100 SNP window, and outlier windows are highlighted. mds = multi-dimensional scaling. **b**, Genomic positions of the haploblocks identified by local PCA across three species of wild sunflowers. Some predicted haploblocks are in multiple pieces due to rearrangements relative to the *H. annuus* reference genome. **c**, PCA for the ann05.01 region in *H. annuus* individuals. **d**, Heterozygosity at ann05.01 for the three clusters of individuals identified in panel c. Box plots show the median, box edges represent the 25^th^ and 75^th^ percentiles, whiskers represent the maximum/minimum data points within 1.5x interquartile range outside box edges. **e**, LD plot for chromosome 5. Upper triangle = all *H. annuus* individuals; lower triangle = only individuals homozygous for the more common haploblock allele. Colours represent the second highest R^2^ value in 0.5 Mbp windows. Purple bars and the white box represent ann05.01. **f**, Comparison of chromosome 5 organization between two reference assemblies of cultivated sunflower. **g**, Comparison between genetic maps for two mapping populations of *H. petiolaris fallax* and the chromosome 17 sequence of the reference HA412-HOv2 sunflower assembly. A predicted haploblock (pet17.01) is highlighted in purple. **h**, Comparison of HiC link patterns between pairs of individuals with different haplotypes at three haploblocks. Red or blue dots show increased long distance interactions in one sample, consistent with structural variants bringing distant regions together. Purple bars and solid black lines represent predicted haploblock regions. The dune haplotype for pet09.01 is split in two fragments (dotted lines) when aligned to the *H. annuus* reference genome.

These patterns match the expectations for large, segregating structural variants (SVs). Theoretical and empirical work indicates that SVs can facilitate adaptive divergence in the face of gene flow by reducing recombination between locally adaptive alleles^29, 30^. In particular, inversions have been shown to control adaptive phenotypic variation (e.g. migration^31^, colour^32^, flowering time^33^), and to be associated with environmental clines^34, 35^. We used three different approaches to determine whether these haploblocks are associated with SVs (Extended Data Table 1). First, we compared the genome assemblies of two cultivars of *H. annuus* that have opposite genotypes at haploblock regions on chromosomes 1 and 5 (ann01.01 and ann05.01). At each of those regions we found one and two large inversions, respectively (Fig 4f; Extended Data Fig. 6a). We also aligned ten *H. annuus* and four *H. petiolaris* genetic maps to the sunflower reference genome; we observed suppressed recombination at ten haploblocks, and evidence for three haploblocks being caused by large inversions (Fig. 4g; Extended Data Fig. 6b,c). Lastly, we used Hi-C sequencing^36^ to compare pairs of early- and late-flowering *H. argophyllus* and dune and non-dune *H. petiolaris*, and looked for differences in physical linkage at haploblock regions. We found support for SVs, ranging from likely full-length inversions to more complex rearrangements, at 11 regions in *H. petiolaris* and one in *H. argophyllus* (Fig. 4h, Extended Data Fig. 7). For one haploblock for each species, we could find no evidence of SVs in our HiC data, suggesting that recombination between haplotypes might be suppressed by other mechanisms in these regions. We also confirmed the presence of large SVs underling four of the haploblocks we detected in wild *H. annuus* by comparing these Hi-C data to those for the HA412-HO reference cultivar (itself *H. annuus*; Extended Data Fig. 7). These results point to SVs being associated with most of the haploblock regions we detected.

Of the 37 haploblocks we identified, two correspond to the chromosome 6 region associated with flowering time in *H. argophyllus* (arg06.01 and arg06.02), and three with the *H. petiolaris* seed size, flowering time and CEC plateaus (pet09.01, pet11.01 and pet14.01; Fig. 4b; Extended Data Table 1). We also identified four additional haploblocks co-localizing with regions of high genetic differentiation between dune and non-dune ecotypes of *H. petiolaris* (Fig. 3f, Extended Data Fig. 3d), bringing the total of dune-associated haploblocks to seven, four of which are shared between both independent dune ecotypes (Texas and Colorado; Extended Data Fig. 5). Phylogenetic analysis finds that these dune-associated haploblocks predate the split between the *H. petiolaris* subspecies *fallax* and *petiolaris*, and that five are polymorphic in both subspecies (Fig. 5b; Extended Data Fig. 3e). Such high levels of divergence are common to most haploblock regions (Fig. 5a). For the two haploblocks that are polymorphic between *H. annuus* reference genomes (ann01.01 and ann05.01; Fig. 4f; Extended Data Fig. 6a), sequence identity between haplotypes is 94-95%, much lower than the 99.4% for the rest of the genome. Divergence times between all but one of the haploblocks exceed 1 MYA, in most cases (32/37) before the *H. annuus-H. argophyllus* speciation event^37^ (Fig. 5b). This seems at odds with the observation that haploblock polymorphisms are not shared between sunflower species. Ancient haploblocks could have been maintained in selected lineages, possibly by balancing selection^38^, but this should result in transpecific polymorphisms. Alternatively, they could be more recently introgressed from divergent taxa, a hypothesis supported for four *H. argophyllus* haploblocks, in which one haplotype is phylogenetically closer to *H. annuus* than to *H. argophyllus* (Fig. 2m). However, a donor species could not be identified for more divergent haploblocks, raising the intriguing possibility that they may be introgressed from one or more extinct taxa.

**Fig. 5:**
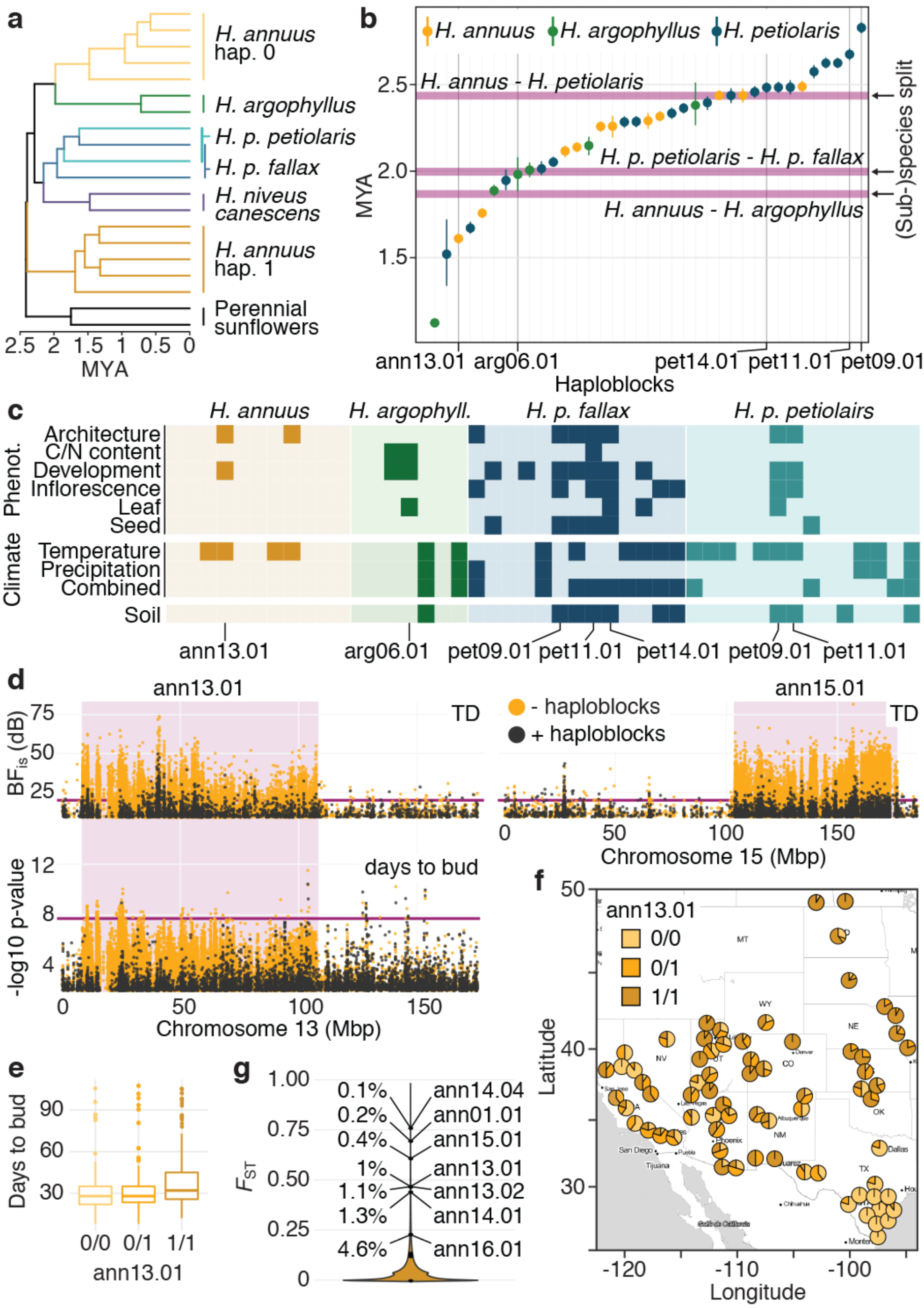
Haploblocks are highly divergent and are associated with multiple ecologically-relevant traits and environmental variables. **a**, Bayesian phylogeny of ann05.01. MYA = million years ago. **b**, Divergence time estimates for haploblocks, relative to those of different (sub-)species of wild sunflowers. **c**, Summary of GWA and GEA analyses for haploblocks treated as single loci. Darker squares represent categories containing at least one trait/variable significantly associated with a haploblock (p < 0.001; BF_is_ > 10). **d**, GEAs for temperature difference (TD), a measure of continentality, and GWA for flowering in *H. annuus*, using a kinship matrix and PCA covariate including (black dots) or excluding (yellow dots) the haploblock regions. Haploblock regions are highlighted in purple. Purple lines represent BF_is_ = 20 dB in GEA plots, and 5% significance after Bonferroni correction in the GWA plot. Only positions with BF_is_ > 9 dB or -log10 p-value > 2 are plotted. **e**, Flowering time for individuals with different genotypes at ann13.01. Box plots show the median, box edges represent the 25^th^ and 75^th^ percentiles, whiskers represent the maximum/minimum data points within 1.5x interquartile range outside box edges. **f**, Geographic distribution of ann13.01 haplotypes. **g**, Distribution of F_ST_ values for individual SNPs and haploblocks in comparisons between *H. annuus texanus* and other *H. annuus* populations. Percentiles are reported for the most highly divergent haploblocks.

## Haploblocks underlie ecotype divergence

As we have shown, haploblocks can have remarkably strong associations with phenotypic traits and environmental variables (Fig. 2c, 3f; Extended Fig. 2b, 3b,c), but these examples represent only a small proportion of the total haploblock regions we identified. Are other haploblocks also involved in local adaptation? Theory suggests that SVs are likely to establish by capturing multiple adaptive alleles^39^; consistent with this, when we treated haploblocks as individual loci, we found that haploblocks are often associated with multiple types of traits (Fig 5c; Extended Data Fig. 8,9).

Surprisingly, some of the strongest association we identified with this approach did not show up in our initial GWA and GEA analyses. Haploblocks are in fact large enough to affect the genome-wide estimates of relatedness between individuals (kinship and PCA) routinely employed to compensate for population structure in GWA and GEA analyses, which can result in their association signal being masked^40^. This is particularly evident for ann13.01, which at ∼100 Mbp is the largest of the haploblocks we identified; significant plateaus for temperature difference (TD, a measure of climate continentality) and flowering time are only revealed once haploblock regions are removed from the kinship covariate (Fig 5d,e). Interestingly, this and several other haploblocks appear to differentiate Texas populations of *H. annuus* from the rest of the range (Fig. 5f; Extended Data Fig. 5), consistent with the distribution of the *texanus* ecotype of *H. annuus*^41^. Similar to the *H. petiolaris* dune comparison (Extended Data Fig. 3d), haploblocks are more differentiated than SNPs in comparisons between Texas and other populations (t(10) = 4.01, p = 0.0024, two-sided T-test; Fig. 5g), supporting a role for haploblocks in local adaptation of this subspecies, or in increasing its reproductive isolation with local congeners (i.e. reinforcement^42^).

## Conclusions

We identified a large number of highly divergent, multi-Mbp-long haploblocks in wild sunflowers, many of which appear to underlie ecotype formation; four in the early flowering ecotype from *H. argophyllus*; seven in the *texanus* ecotype of *H. annuus*; and seven in dune ecotypes of *H. petiolaris* (Extended Data Fig. 5). These haploblocks are often, but not always, linked to large SVs (especially inversions), which provide a straightforward mechanism for suppressing recombination between haplotypes, therefore maintaining adaptive allelic combinations. The total number and effects of such haplotypes are likely even larger than this, since our approach is biased towards detection of divergent and large (>1 Mbp) haploblocks. Ecotypic differentiation is often seen as a first step toward the generation of new species^43^, and the ecotypes discussed above appear to represent different stages in the speciation continuum. The coastal island ecotype of *H. argophyllus* is least divergent, and the only reproductive barrier we are aware of is the difference in flowering time^7^, which provides only modest protection from gene flow. In contrast, multiple intrinsic and extrinsic reproductive barriers differentiate the two dune ecotypes of *H. petiolaris fallax* from nearby non-dune populations^6, 44^, reducing but not eliminating gene migration^16, 45^. It is noteworthy that several haploblocks are associated with both traits favouring local adaptation and contributing to reproductive isolation (e.g., seed size and flowering time, respectively, in the dune ecotypes), an architecture that facilitates speciation with gene flow^29^. More generally, flowering time mapped to one or more haploblocks in all ecotypes, suggesting that it plays an especially important role in successful ecotype formation, perhaps due to its dual role in local adaptation and assortative mating^46^.

An unanswered question is how the linked combinations of locally favoured mutations found in haploblocks arose. Possibly, sets of locally adaptive alleles initially developed in geographically isolated populations^47^. Secondary contact and hybridization would favour the evolution of reduced recombination among such alleles through the establishment of SVs^39^ or other recombination modifiers^28^. An origin through introgression would also help account for the high divergence and massive size of many of the haploblocks, as well as the lack of shared haploblock polymorphisms between species. The alternative explanation for the latter, incomplete lineage sorting, would require extensive haploblock loss. After haploblock establishment (regardless of how it occurred), new locally adaptive mutations would be more likely to persist under migration-selection balance if linked to other adaptive alleles^48, 49^, potentially leading to the outsized effects reported here. Our work reveals a modular genetic architecture underlying ecotype formation, a surprising and unforeseen origin of many locally adapted gene modules though introgression, and a critical role of recombination modifiers, especially structural variants, in adaptive divergence with gene flow.

## Methods

### Seed and soil collection

During the summer of 2015 we visited 192 wild populations spanning the native distribution of *H. annuus, H. petiolaris,* and *H. argophyllus*, and collected seeds from 21-37 individuals from each population. Seeds from ten additional populations of *H. annuus* had been previously collected in the summer of 2011. Three to five soil samples (0 - 25 cm depth) were collected with a corer at each population, from across the area in which seeds were collected. Soils were air dried in the field, further dried at 60 °C in to the lab, and passed through a 2 mm sieve to remove roots and rocks. Soils were then submitted to Midwest Laboratories Inc. (Omaha, NE, USA) for analysis.

### Common garden

Ten plants from each of 151 selected populations were grown at the Totem Plant Science Field Station of the University of British Columbia (Vancouver, Canada) in the summer of 2016. Pairs of plants from the same population of origin were sown using a completely randomized design. At least three flowers from each plant were bagged before anthesis to prevent pollination, and manually crossed to an individual from the same population of origin. Phenotypic measurements were performed throughout plant growth, and leaves, stem, inflorescences and seeds were collected and digitally imaged to extract relevant morphometric data (see Supplementary Table 1).

### Library preparation and sequencing

Whole-genome shotgun (WGS) sequencing libraries were prepared for 719 *H. annuus*, 488 *H. petiolaris*, 299 *H. argophyllus* individuals, and twelve additional samples from annual and perennial sunflowers (Supplementary Table 1). Genomic DNA was sheared to ∼400 bp fragments using a Covaris M220 ultrasonicator (Covaris, Woburn, Massachusetts, USA) and libraries were prepared using a protocol largely based on Rowan *et al.*, 2015^51^, the TruSeq DNA Sample Preparation Guide from Illumina (Illumina, San Diego, CA, USA) and Rolhand *et al.*, 2012^52^. In order to reduce the proportion of repetitive sequences, libraries were treated with a Duplex-Specific Nuclease (DSN; Evrogen, Moscow, Russia), following the protocols reported in Shagina *et al.* 2010^10^ and Matvienko *et al.* 2013^53^, with modifications (see Supplementary Methods for details). All libraries were sequenced at the McGill University and Génome Québec Innovation Center on HiSeq2500, HiSeq4000 and HiSeqX instruments (Illumina, San Diego, CA, USA), to produce paired end, 150 bp reads. Libraries with fewer reads were re-sequenced to increase genome coverage. After quality filtering (see below), a total of 60.7 billion read pairs were retained, equivalent to 14.5 Tbp of sequence data.

### Variant calling

The call set included the 1518 samples described above, the Sunflower Association Mapping (SAM) population (a set of cultivated *H. annuus* lines ^54^), and wild *Helianthus* samples previously sequenced for other projects^54–56^, for a total of 2392 samples (Supplementary Table 1). The additional samples were included to improve SNP calling, and to identify haploblock genotypes. Sequences were trimmed for low quality using Trimmomatic^57^ (v0.36) and aligned to the *H. annuus* XRQv1 genome^9^ using NextGenMap^58^ (v0.5.3). We followed the best practices recommendations of The Genome Analysis ToolKit (GATK)^59^, and executed steps documented in GATK’s germline short variant discovery pipeline (for GATK 4.0.1.2). During genotyping, to reduce computational time and improve variant quality, genomic regions containing transposable elements were excluded^9^. Since performing joint-genotyping on the whole ensemble of samples would have been computationally impractical, genotyping was performed independently on three per-species cohorts (*H. annuus*, *H. argophyllus* and *H. petiolaris*).

### Variant quality filtering

Genotyping produced VCF files featuring an extremely large number of variant sites (222M, 78M and 167M SNPs and indels for *H. annuus, H. argophyllus* and *H. petiolaris*, respectively). Over the called portion of the genome, this corresponds to 0.07 to 0.2 variants per bp, with 30-47% percent of variable sites being indel variation. To remove low-quality calls and produce a dataset of a more manageable size, we used GATK’s VariantRecalibrator (v4.0.1.2), which filters variants in the call set according to a machine learning model inferred from a small set of “true” variants. In the absence of an externally-validated set of known sunflower variants to use as calibration, we computed a stringently-filtered set from top-N samples with highest sequencing coverage for each species (N=67 (SAM) samples for *H. annuus*, and N=20 otherwise). The stringency of the algorithm in classifying true/false variants was adjusted by comparing variant sets produced for different parameter values (tranche 100.0, 99.0, 90.0, 70.0, and 50.0). For each cohort, results for tranche = 90.0 were chosen for downstream analysis, based on heuristics: the number of novel SNPs identified, and improvements to the transition/transversion ratio (towards GATK’s default target of 2.15).

### Remapping sites to the HA412-HOv2 reference genome

Our initial analysis of haploblocks (see section “Population genomic detection of haploblocks”), as well as GWA/GEA results for haploblocks regions, found many instances of disconnected haploblocks and high linkage between distant parts of the genome, suggesting problems in contig ordering. We remapped genomic locations from XRQv1^9^ to HA412-HOv2^11^ using BWA^60^. Measures of LD using vcftools^61^ showed that remapping significantly improved LD decay (Extended Data Fig. 1a) and produced more contiguous haploblocks (Extended Data Fig. 1b), supporting the accuracy of the new genome assembly and our remapping procedure. While we recognize that this approach reduces accuracy at the local scale, and would not be appropriate, for example, for determining the effects of variants on coding sequences, it produces a more accurate reflection of the genome and linkage structure.

### Phylogenetic analysis

Variants were called for 20 windows of 1 Mbp, randomly selected across the genome. Indels were removed and SNP sites were filtered for <20% missing data and minor allele frequency >0.1%. All sites were then concatenated and analyzed using IQtree^62–64^ with ascertainment bias and otherwise default parameters.

### Genome-wide association mapping

Genome-wide association analyses were performed for 86, 30 and 69 phenotypic traits in *H. annuus, H. argophyllus* and *H. petiolaris,* respectively, using the EMMAX (v07Mar2010) or the EMMAX module in EasyGWAS^65^; an annotated list of candidate genes is reported in Supplementary Table 2. Inflorescence and seed traits could not be collected for *H. argophyllus*, since most plants of this species flowered very late in our common garden, and failed to form fully-developed inflorescences and set seeds before temperatures became too low for their survival.

### Genome-environment association analyses

Twenty-four topo-climatic factors were extracted from climate data collected over a 30-year period (1961-1990) for the geographic coordinates of the population collection sites, using the software package Climate NA^66^. Soil samples from each population were also analyzed for 15 soil properties (Supplementary Table 1). The effects of each environmental variable were analyzed using BayPass^67^ version 2.1. Following Gautier, 2015^67^, we employed Jeffreys’ rule^68^, and quantified the strength of associations between SNPs and variables as “strong” (10 dB ≤ BF_is_ < 15 dB), “very strong” (15 dB ≤ BF_is_ < 20 dB) and decisive (BF_is_ ≥ 20 dB). An annotated list of candidates genes from GEA analyses is reported in Supplementary Table 2.

### Transgenes and expression assays

The complete coding sequences (CDS) of *HaFT1*, *HaFT2* and *HaFT6* were amplified from complementary DNA (cDNA) from *H. argophyllus* individuals carrying the early and late haplotype for arg06.01. Two alleles of the *HaFT2* CDS were identified in late-flowering *H. argophyllus* plants (one of them identical to the *HaFT2* CDS from early-flowering individuals), differing only for two synonymous substitutions at position 285 and 288. All alleles were placed under control of the constitutive CaMV 35S promoter in pFK210 derived from pGREEN^69^. Constructs were introduced into plants by *Agrobacterium tumefaciens*-mediated transformation^70^. Col-0 and *ft*-10 seeds were obtained from the Arabidopsis Biological Resource Center. All primer sequences are reported in Supplementary Table 3.

### Population genomic detection of haploblocks

The program lostruct (local PCA/population structure) was used to detect genomic regions with abnormal population structure^28^. Lostruct divides the genome into non-overlapping windows and calculates a PCA for each window. It then compares the PCAs derived from each window and calculates a similarity score. The matrix of similarity scores is then visualized using a multidimensional scaling (MDS) transformation. Lostruct analyses were performed on the *H. annuus*, *H. argophyllus*, *H. petiolaris petiolaris*, and *H. petiolaris fallax* datasets, as well as in a *H. petiolaris* dataset including both *H. petiolaris petiolaris* and *H. petiolaris fallax* individuals. For each dataset, lostruct was run with 100 SNP-wide windows and independently for each chromosome. Each MDS axis was then visualized by plotting the MDS score against the position of each window in the chromosome.

Many localized regions of extreme MDS values with high variation in MDS scores and sharp boundaries were detected (Fig. 4a; Extended Data Fig. 4). Localized changes to population structure could occur due to selection or introgression, but both the size and discrete nature of the regions are consistent with underlying structural changes defining the boundaries and preventing recombination. For example, inversions prevent recombination between orientations and if inversion haplotypes are diverged enough, they will show up in lostruct scans^28^. Since we are interested in recombination suppression in the context of adaptation, we focused on regions that had the following features: (1) a PCA in the region should divide samples into three groups representing 0/0, 0/1 and 1/1 genotypes, (2) the middle 0/1 genotype should have higher average heterozygosity and (3) there should be high linkage disequilibrium (LD) within the region.

The combined evidence of PCA and linkage suggests that the lostruct outlier regions are characterized by long haplotypes with little or no recombination between haplotypes. We refer to these as haploblocks. To explore the haplotype structure underlying the haploblocks, sites correlated (*R^2^* > 0.8) with PC1 in the PCA of the haploblock were extracted as haplotype diagnostic sites and used to genotype the haploblocks. Since there is seemingly little recombination between haplotypes, this is conceptually similar to a hybrid index and we expect all samples to be consistently homozygous for one haplotypes alleles or be heterozygous at all sites (i.e. similar to an F_1_ hybrid). Haploblock genotypes were assigned to all samples using equation (1), where *p* is the proportion of haplotype 1 alleles and *h* is the observed heterozygosity. The haplotype structure was also visualized by plotting diagnostic SNP genotypes for each sample, with samples ordered by the proportion of alleles from haplotype 1 (e.g. Fig. 2f).

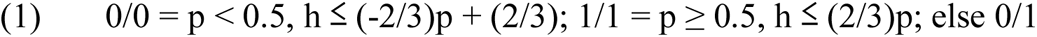

Lostruct was run in SNP datasets containing *H. petiolaris petiolaris, H. petiolaris fallax,* and both subspecies together. Although each dataset produced a collection of haploblocks, they were not identical. Some haploblocks were identified in one subspecies, but not the other, and some were only identified when both subspecies were analyzed together. In some cases, it was clear that haploblocks identified in both subspecies represented the same underlying haploblock because they physically overlapped and had overlapping diagnostic markers. We manually curated the list of haploblocks and merged those found in multiple datasets. We set the boundaries of these merged haploblocks to be inclusive (i.e. include windows found in either) and the diagnostic markers to be exclusive (i.e. only include sites found in both). For this merged set of haploblocks, all *H. petiolaris* samples were genotyped using diagnostic markers.

### Design of genetic markers for haploblock screening

Diagnostic SNPs for haploblocks were extracted from filtered vcf files. The resulting markers Cleaved-Amplified Polymorphic Sequence (CAPS) or direct sequencing markers were tested on representative subsets of individuals included in the original local PCA analysis (Fig. 4a, Extended Data Fig. 4), for which the genotype at haploblocks of interest was known. Marker information are reported in Supplementary Table 3.

### Sequencing coverage analysis

To detect the presence of potential deletions in the late-flowering allele of arg06.01, SNP in the haploblock region with average coverage of at least 4 across at least one of the genotypic classes were selected (in order to exclude positions with overall low mapping quality). SNP positions with coverage 0 or 1 in one genotypic class were counted as missing data for that genotypic class (Extended Data Fig. 2c).

### *H. annuus* reference assemblies comparisons

Masked reference sequences for the *H. annuus* cultivars HA412-HOv2 and PSC8^11, 12^ were aligned using MUMmer^71^ (v4.0.0b2). The programs nucmer (parameters -b 1000 -c 200 -g 500) and dnadiff within the MUMmer package were used. Only orthologous chromosomes were aligned together because of the high similarity and known conservation of chromosome structure. The one-to-one output file was then visualized in R and only included alignments where both sequences were > 5000 bp. Inversion boundaries and sequence identity between haplotypes were further determined using Syri^72^.

### Genetic maps comparisons

Fourteen genetic maps were used: the seven *H. annuus* genetic maps used in the creation of the XRQv1 genome^9^; three newly generated *H. annuus* maps obtained from wild X cultivar F_2_ populations (E.B.M.D., M.T., G.L.O., L.H.R., in preparation); two previously published *H. petiolaris* genetic maps obtained from F_1_ crosses^50^; and two newly generated *H. petiolaris* maps (K.H., Rose L. Andrews, G.L.O., K.L.O., L.H.R., in preparation). Whenever necessary, marker positions relative to XRQv1 were re-mapped to the HA12-HOv2 assembly (see above). Six of the previously described *H. annuus* maps were obtained from crosses between cultivars (the seventh one was obtained from a wild X cultivar cross); in order to determine which haploblock could be expected to segregate in the genetic maps, all of the *H. annuus* SAM population lines were genotyped for each *H. annuus* haploblock using diagnostic markers identified in wild *H. annuus*. Ann01.01 and ann05.01 were found to be highly polymorphic in the SAM population, while other haploblocks were fixed or nearly fixed for a single allele. For all fourteen maps, marker order was compared to physical positions in the HA412-HOv2 reference assembly, and evidence for suppressed recombination or structural variation was recorded (Extended Data Table 1).

### Hi-C

Pairs of *H. petiolaris* and *H. argophyllus* populations that diverged for a large number of haploblocks were selected. Individuals from these populations were genotypes using haploblock diagnostic markers (see “Design of genetic markers for haploblock screening”) to identify, for each species, a pair of individuals with different genotypes at the largest possible number of haploblocks. Chromosome conformation capture sequencing^36, 73^ (Hi-C) libraries were prepared by Dovetail Genomics (Scotts Valley, CA, USA) and sequenced on a single lane of HiSeq X with 150 bp paired end reads. Reads were trimmed for enzyme cut site and base quality using the tool *trim* in the package HOMER^74^ (v4.10) and aligned to the HA412-HOv2 reference genome using NextGenMap^58^ (v0.5.4). Interactions were quantified using the calls ‘makeTagDirectory - tbp 1-mapq 10’ and ‘analyzeHiC -res 1000000 -coverageNorm’ from HOMER. Hi-C data were used in two ways to identify structural changes. First, the difference between interaction matrices for samples of the same species was plotted for each haploblock region where the two samples had different genotypes. Second, the difference between interaction matrices for *H. annuus* (using the HiC data that were generated to scaffold the HA412-HOv2 reference assembly^11^) and each *H. petiolaris* and *H. argophyllus* sample were plotted.

### Haploblock phenotype and environment associations

Since haploblocks are large enough to affect genome wide population structure, their associations with phenotypes of environmental variables may be masked when controlling for population structure. Therefore, a version of the variant file was created with all haploblock sites removed; GWA and GEA analyses were performed as before, but kinship, PCA and genetic covariance matrix were calculated using this haploblock-free variant file. Regions of high associations co-localizing with haploblock regions were identified, and haploblocks were also directly tested by coding each haploblock as a single bi-allelic locus.

To examine the relative importance of haploblocks to trait evolution and environmental adaptation, association results were compared between haploblocks and SNPs. Using SNPs as a baseline allows to control for the correlation between traits or environmental variables. To make values comparable, both SNPs and SVs with minor allele frequency ≤ 0.03 were removed. Each locus was classified as associated (p < 0.001 or BF_is_ > 10 dB) or not to each trait. The number of traits or climate variable each locus was associated with was then counted. The proportion of loci with ≥ 1 traits/climate variables associated for SNPs and haploblocks was then compared using prop.test in R^75^ (Extended Data Fig. 9b).

### Haploblocks phylogenies and dating

The phylogeny of each haploblock region was estimated by Bayesian inference using BEAST^76^ 1.10.4 for 100 genes within the region. The dataset was partitioned, assuming unlinked substitution and clock models for the genes, and analyzed under the HKY model with 4 Gamma categories for site heterogeneity: a strict clock, a “Constant Size” tree prior with a Gamma distribution with shape parameter 10.0 and a scale parameter 0.004 for the population size. Default priors were used for the other parameters. A custom Perl script was used to combine FASTA sequences and the model parameters into XML format for BEAST input. The Markov chain Monte Carlo (MCMC) process was run for 1 million iterations and sampled every 1000 states. The convergence of chains was inspected in Tracer^77^ 1.7.1. In order to estimate divergence times, the resulting trees were calibrated using a mutation rate estimate of 6.9 × 10^−9^ substitutions/site/year for sunflowers^78^, and visualized with R package ggtree^79^ and Figtree v1.4.4^80^. Divergence times were extracted from the trees and plotted showing the 95% highest posterior density (HPD) interval based on the BEAST posterior distribution. This was repeated for 100 non-haploblock genes to estimate the species divergence times.

For the 10 Mb region on chromosome 6 controlling flowering time in *H. argophyllus*, the early flowering haplotype grouped with *H. annuus*. To determine if it is the product of an ancient haplotype that has retained polymorphism only in *H. annuus* or if it is introgressed from *H. annuus*, the phylogeny of 10 representative *H. argophyllus* samples homozygous for each haploblock allele, as well as 200 *H. annuus* samples, was inferred using IQtree. SNPs from the 10 Mb region were concatenated and the maximum likelihood tree was constructed using the GTR model with ascertainment bias correction. Branch support was estimated using ultrafast bootstrap implemented in IQtree^62–64^ with 1,000 bootstrap replicates. Phylogenies of haploblock arg03.01, arg03.02 and arg06.02 were inferred using the same approach. To explore intra-specific history of the *H. petiolaris* haploblocks, all samples homozygous for either allele for each haploblock were selected, and phylogenies were constructed using IQtree with the same settings.

### Data availability

All raw sequenced data are stored in the Sequence Read Archive (SRA) under BioProjects PRJNA532579, PRJNA398560 and PRJNA564337. Accession numbers for individual samples are listed in Supplementary Table 1. The HA412-HOv2 and PSC8 genome assemblies are available at https://sunflowergenome.org/ and https://heliagene.org/. GWA results are available at https://easygwas.ethz.ch/gwas/results/xxx/. *HaFT1*, *HaFT2* and *HaFT6* sequences have been deposited in GenBank under accession numbers MN517758-MN517761.

### Code availability

All code associated with this project is available at https://github.com/owensgl/haploblocks.

## Acknowledgments

We thank Jérôme Gouzy and Nicolas B. Langlade for providing access to the HA412-HOv2 annotation and PSC8 genome assembly, Brook T. Moyers for discussion and providing the *H. argophyllus* picture, Julie Lee-Yaw and Armando J. Moreno-Geraldes for comments, Dominique Skonieczny, Amy Kim, Ana Parra and Cassandra Konecny for assistance with field work and data acquisition, James D. Herndon for providing the dune *H. petiolaris* picture, Mihir Nanavati and Andrew Warfield for computing advice, and Compute Canada for computing resources. Funding was provided by Genome Canada and Genome BC (LSARP2014-223SUN), the NSF Plant Genome Program (DBI-1444522), the International Consortium for Sunflower Genomic Resources, a HFSP long-term postdoctoral fellowship to M.T. (LT000780/2013), a Banting postdoctoral fellowship to G.L.O.

## Author Contributions

L.H.R., S.Y., J.M.B, L.A.D., N.B. conceived the study; M.T., N.B., D.O.B., E.B.M.D., I.I., M.A.P., W.C., L.A.D. collected data and performed experiments; M.T., G.L.O., J.S.L., S.S., K.H., K.L.O., E.B.M.D., K.L. analyzed data; S.E.S., S.M. contributed resources; R.N. provided conceptual advice; M.T., G.L.O., L.H.R. wrote the manuscript, with contributions from all the authors.

## Additional Information

**Supplementary Information** is available for this paper.

## Extended Data

**Extended Data Table 1:** Positions and frequencies of haploblocks, and experimental support for linked SVs. Allele frequencies for haplotype 1 are reported. For *H. petiolaris*, allele frequencies for *H. p. fallax* and *H. p. petiolaris*, respectively, are reported. Hi-C geno = haploblock genotypes of the pair of individuals that were used for Hi-C sequencing (for *H. annuus* haploblocks, only the genotype of HA412-HO is reported). Experimental support for haploblock: H = differences in Hi-C patterns between samples; h = differences in Hi-C patterns relative to the HA412-HO reference; A = differences between reference *H. annuus* assemblies; G = differences in orientation between genetic maps; g = recombination suppression in genetic maps; No SV = no evidence of SV in Hi-C experiments despite appropriate comparison. Haploblock positions are relative to the HA412-HOv2 assembly; in some cases predicted haploblocks are in multiple pieces due to rearrangements relative to the reference.

| Species | Haploblock ID | Chr. | Allele freq. | Hi-C geno. | Support | Size (Mbp) | Region (Mbp) |
| --- | --- | --- | --- | --- | --- | --- | --- |
| *H. annuus* | ann01.01 | 1 | 0.21 | 0/0 | A, h | 4 | 6-10 |
| *H. annuus* | ann05.01 | 5 | 0.52 | 0/0 | A, g | 29 | 148-177 |
| *H. annuus* | ann11.01 | 11 | 0.25 | 0/0 | g | 35 | 19-54 |
| *H. annuus* | ann13.01 | 13 | 0.62 | 1/1 | h, g | 101 | 9-110 |
| *H. annuus* | ann13.02 | 13 | 0.13 | 0/0 | g | 19 | 139-158 |
| *H. annuus* | ann14.01 | 14 | 0.22 | 0/0 |  | 28 | 101-129 |
| *H. annuus* | ann14.02 | 14 | 0.9 | 1/1 |  | 6 | 129-135 |
| *H. annuus* | ann15.01 | 15 | 0.31 | 0/0 | h, g | 72 | 104-176 |
| *H. annuus* | ann16.01 | 16 | 0.85 | 1/1 |  | 13 | 10-23 |
| *H. annuus* | ann16.02 | 16 | 0.96 | 1/1 | h, g | 12 | 147-159 |
| *H. annuus* | ann17.01 | 17 | 0.81 | 1/1 |  | 9 | 188-197 |
| *H. argophyllus* | arg03.01 | 3 | 0.89 | 1/1 : 1/1 |  | 47 | 42-89 |
| *H. argophyllus* | arg03.02 | 3 | 0.06 | 0/0 : 0/0 |  | 14 | 28-42 |
| *H. argophyllus* | arg06.01 | 6 | 0.08 | 1/1 : 0/0 | No SV | 25 | 130-155 |
| *H. argophyllus* | arg06.02 | 6 | 0.05 | 0/0 : 0/0 |  | 5 | 125-130 |
| *H. argophyllus* | arg10.01 | 10 | 0.19 | 0/1 : 0/0 | H | 29 | 161-190 |
| *H. argophyllus* | arg16.01 | 16 | 0.14 | 0/0 : 0/0 |  | 3 | 202-205 |
| *H. petiolaris* | pet01.01 | 1 | 0.67; 0.97 | 1/1 : 0/0 | H, g | 20 | 0-12, 91-99 |
| *H. petiolaris* | pet05.01 | 5 | 0.91; 0.62 | 0/0 : 1/1 | H, G, g | 28 | 154-186 |
| *H. petiolaris* | pet06.01 | 6 | 0.23; 0 | 0/0 : 0/1 | H | 23 | 56-79 |
| *H. petiolaris* | pet06.02 | 6 | 0; 0.11 | 0/0 : 0/0 | h | 37 | 19-56 |
| *H. petiolaris* | pet06.03 | 6 | 0; 0.16 | 0/0 : 0/0 |  | 2 | 9-10 |
| *H. petiolaris* | pet07.01 | 7 | 0.59; 0.08 | 0/0 : 1/1 |  | 14 | 116-130 |
| *H. petiolaris* | pet08.01 | 8 | 0.70; 0.07 | 0/0 : 1/1 | H | 7 | 87-94 |
| *H. petiolaris* | pet09.01 | 9 | 0.17; 0.36 | 1/1 : 0/0 | H, g | 32 | 105-123, 128-141 |
| *H. petiolaris* | pet10.01 | 10 | 0.86; 1 | 0/0 : 1/1 | No SV | 14 | 0-14 |
| *H. petiolaris* | pet11.01 | 11 | 0.78; 0.39 | 0/0 : 1/1 | H, G, g | 62 | 3-65 |
| *H. petiolaris* | pet12.01 | 12 | 0; 0.09 | 0/0 : 0/0 | g | 26 | 89-95, 100-111, 155-164 |
| *H. petiolaris* | pet13.01 | 13 | 0.99; 0.65 | 1/1 : 1/1 |  | 1 | 148-149 |
| *H. petiolaris* | pet14.01 | 14 | 0.88; 0.99 | 0/0 : 1/1 | H | 73 | 98-171 |
| *H. petiolaris* | pet14.02 | 14 | 0.01; 0.68 | 0/0 : 0/0 |  | 9 | 135-144 |
| *H. petiolaris* | pet16.01 | 16 | 0.74; 0.36 | 0/0 : 1/1 | H | 4 | 6-10 |
| *H. petiolaris* | pet16.02 | 16 | 0.79; 1 | 0/0 : 1/1 | H | 53 | 6-23, 34-46, 109-112, 118-123, 137-149, 206-210, 214-218 |
| *H. petiolaris* | pet17.01 | 17 | 0.62; 0.16 | 0/0 : 1/1 | H, G | 11 | 12-23 |
| *H. petiolaris* | pet17.02 | 17 | 1; 0.91 | 1/1 : 1/1 |  | 29 | 39-68 |
| *H. petiolaris* | pet17.03 | 17 | 0.29; 0 | 1/1 : 0/0 | H | 17 | 188-205 |
| *H. petiolaris* | pet17.04 | 17 | 1; 0.90 | 1/1 : 1/1 |  | 11 | 194-205 |

**Extended Data Fig. 1:**
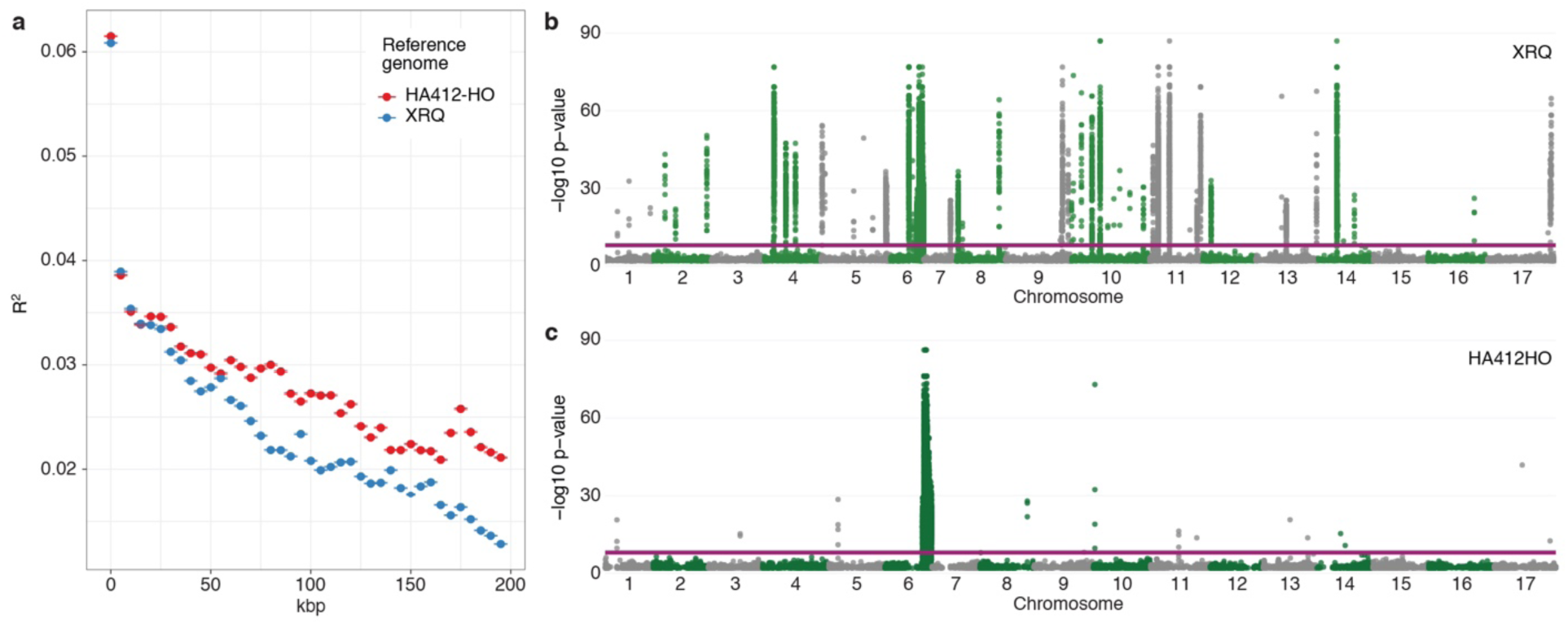
Re-mapping SNPs to the HA412-HOv2 reference genome improved ordering. **a**, Comparison between the original order of SNPs in chromosome 2 on the XRQv1 assembly^9^ (against which sequencing reads were originally mapped) and after SNP re-mapping to the HA412-HOv2 assembly^11^. Data is summarized in 5 kbp ranges. Error bars represent 2 standard errors. The higher R^2^ at longer distances is due to better scaffolding of contigs in HA412-HOv2. **b**, GWA for flowering time in *H. argophyllus* based on the XRQv1 assembly identified more than 40 highly significant associations. **c**, Re-mapping of the SNPs to the new HA412-HOv2 sunflower assembly considerably reduced the number of associations in the flowering time GWA, with the vast majority of the signal mapping to the arg06.01 haploblock region (see Fig. 2). The purple lines represent 5% significance after Bonferroni correction. Only positions with - log10 p-value > 2 are plotted.

**Extended Data Fig. 2:**
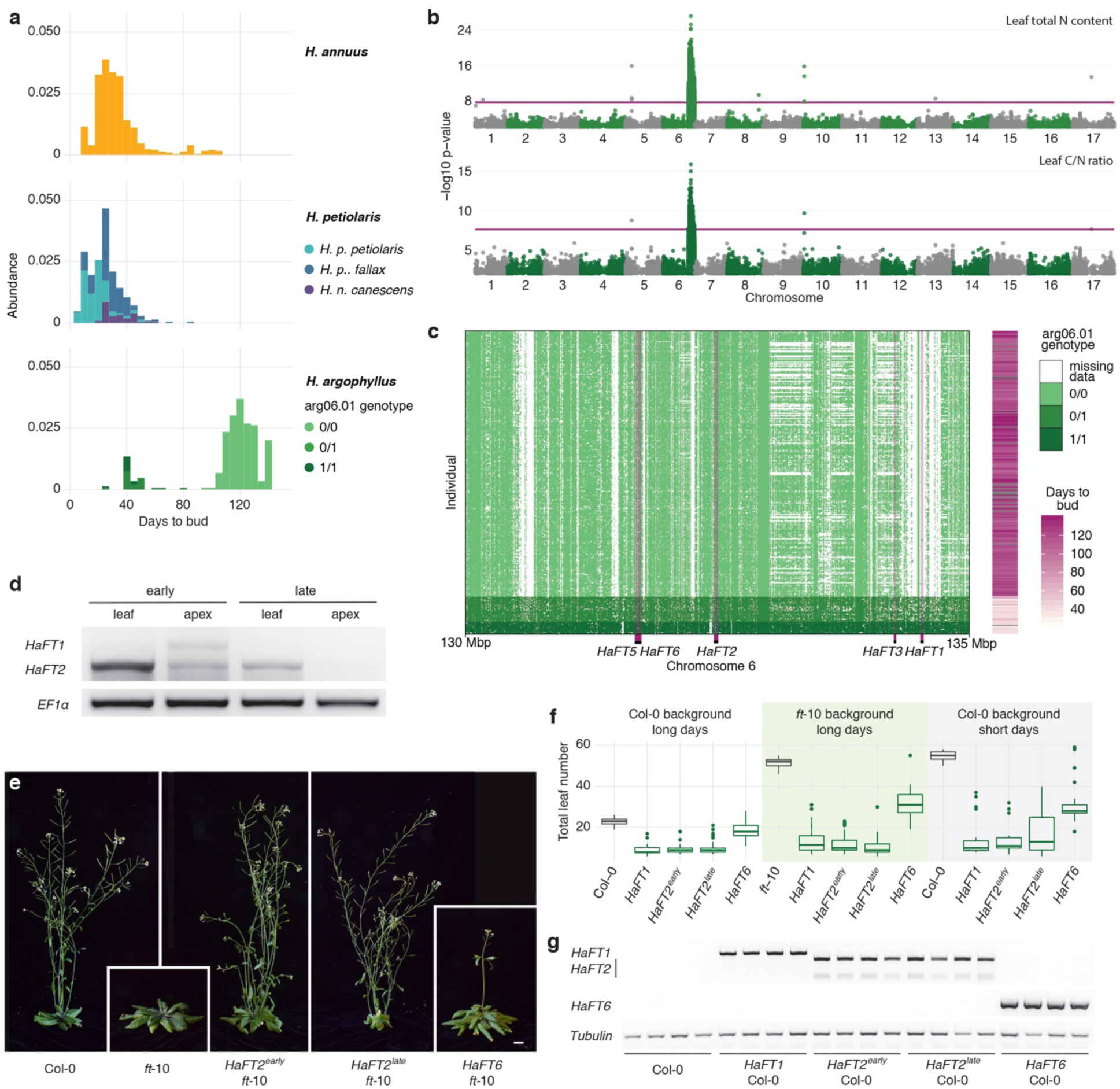
Phenotypic, structural and functional analyses for arg06.01. **a**, Flowering time for the three wild sunflower species measured in a common garden experiment. **b**, Carbon and nitrogen content GWAs in *H. argophyllus.* The purple lines represent 5% significance after Bonferroni correction. Only positions with -log10 p-value > 2 are plotted. **c**, Genotype presence/absence for the 130-135 Mbp region of chromosome 6 in *H. argophyllus*. The x-axis represents consecutive SNP positions; distances on this axis are therefore not proportional to physical distances on the chromosome. Purple bars highlight the positions of the five *HaFT* genes in the region (*HaFT5* and *HaFT6* are only a few hundred bp apart). Flowering time data are the same used in GWA analyses. **d**, *HaFT1* and *HaFT2* expression levels in mature leaves or shoot apices of >6-month-old, flowering *H. argophyllus* plants, grown in a greenhouse in long days conditions (14 hours light; 10 hours dark). This experiment was performed on two independent pairs of individuals, with similar risults. **e**, Six-week-old *A. thaliana* plants grown in long day conditions at 23 °C. **f**, Flowering time in long and short days (10 hours light, 14 hours dark). *HaFT2* alleles from early- and late-flowering *H. argophyllus* plants complement the *ft*-10 mutant, similar to *HaFT1* from the early-flowering ecotype. *HaFT6* is expressed at low levels in *H. argophyllus* plants (not shown), and appears to be a hypo-functional *FT* homolog. Box plots show the median, box edges represent the 25^th^ and 75^th^ percentiles, whiskers represent the maximum/minimum data points within 1.5x interquartile range outside box edges. Differences in flowering time between untransformed controls, *HaFT6* lines and all the other transgenic lines are significant in all conditions (p < 10^-6^; one-way ANOVA with post-hoc Tukey HSD test, df = 4; n ≥ 10). Size bar = 1 cm. **g**, PCR detection of transgene expression in leaves of four independent primary transformants for each transgenic line and wild-type Col-0 plants, grown for four weeks in long days. The reduced ability of *HaFT6* to induce flowering is not due to inefficient expression of the transgene.

**Extended Data Fig. 3:**
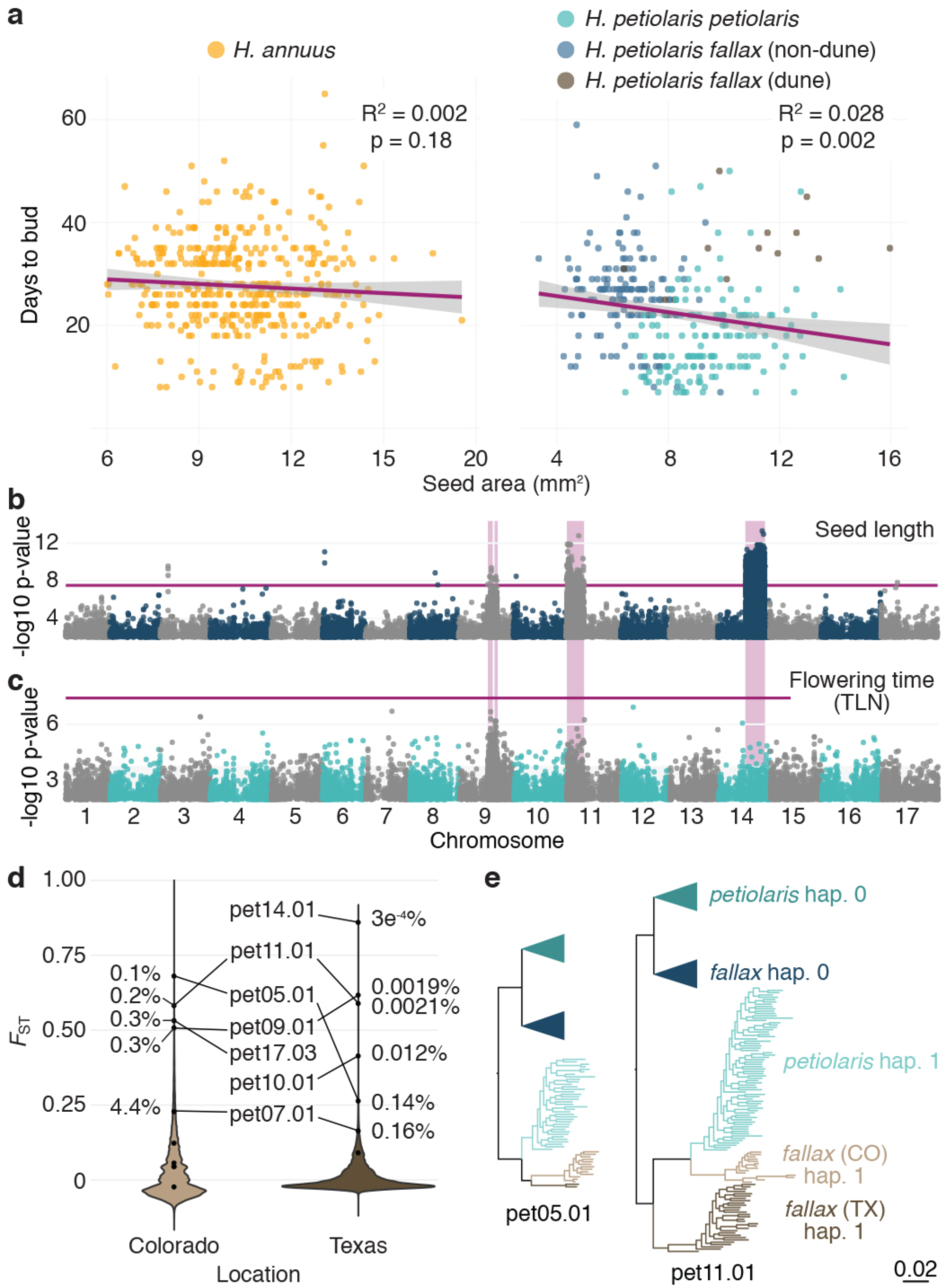
Several haploblocks differentiate dune and non-dune populations of *H. petiolaris*. **a**, Correlation between seed size and flowering time. While dune-adapted *H. petiolaris fallax* flowers later and has larger seeds than non-dune-adapted populations, these two traits generally show no correlation, or a weak negative correlation, in *H. annuus* and *H. petiolaris*. Purple lines represent linear regressions, shaded grey area are 95% confidence intervals. **b**, Seed length GWA in *H. petiolaris fallax*. No significant association with haploblocks is found in GWA analyses for seed width (not shown). **c**, Flowering time (approximated as TLN, total leaf number on the primary stem) GWA for *H. petiolaris petiolaris.* The purple lines in panels c, d represent 5% significance after Bonferroni correction. Only positions with -log10 p-value > 2 are plotted. **d**, Distribution of F_ST_ values for SNPs and haploblocks in comparisons between dune- and non dune-adapted populations of *H. petiolaris fallax* in Colorado and Texas. Percentiles are reported for the most highly divergent haploblocks. **e**, Maximum-likelihood trees for two of the haploblocks segregating within *H. petiolaris*. Dune populations of *H. petiolaris fallax* are highlighted in light (Colorado) and dark tan (Texas). For pet09.01 and pet11.01, although both dune populations have converged on the same haplotype, the Texas haplotype is the ancestral *H. petiolaris fallax* copy, while in Colorado the haplotype is derived from introgression with *H. petiolaris petiolaris*, suggesting convergent adaptation.

**Extended Data Fig. 4:**
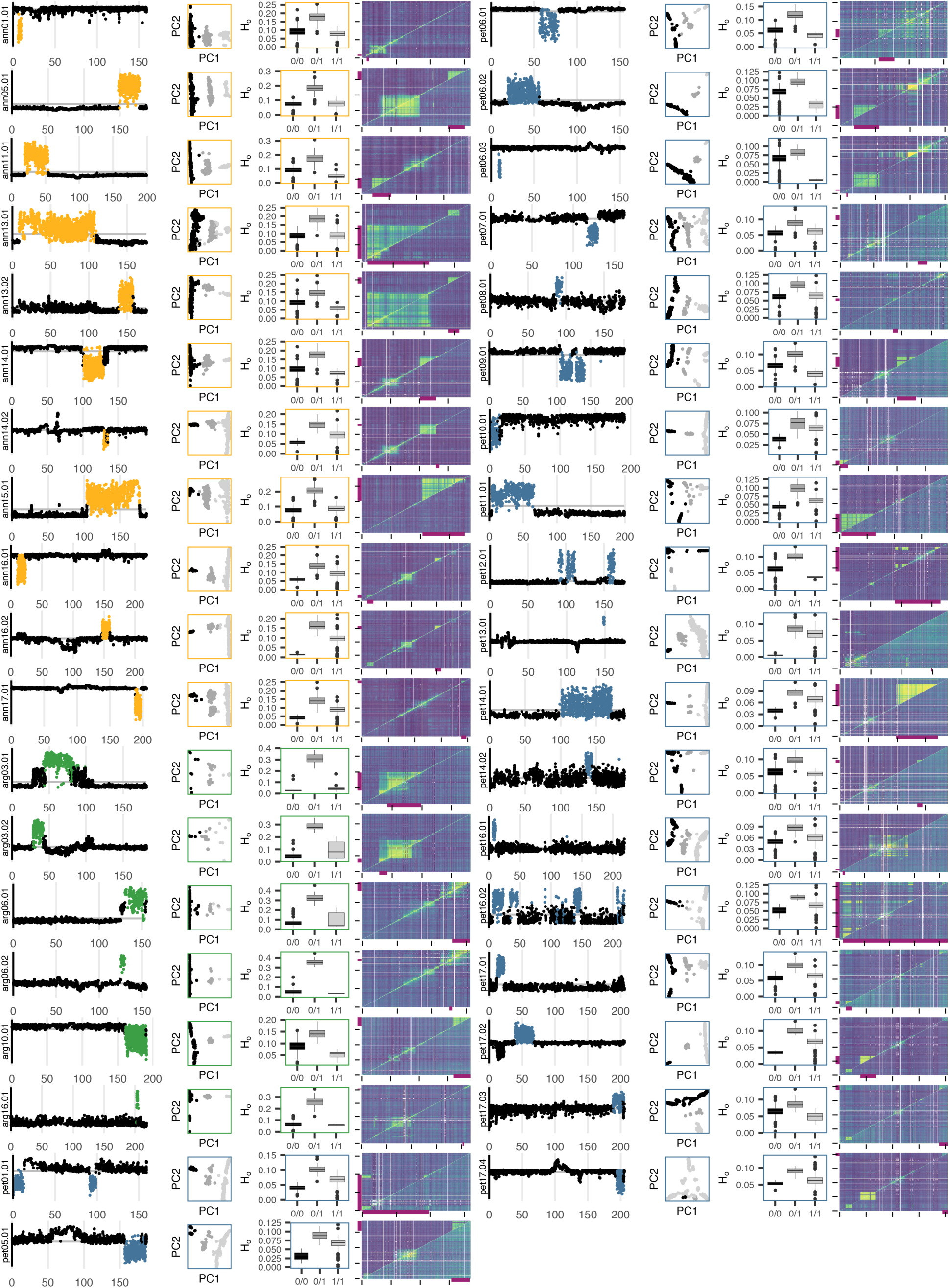
Local PCA highlights haploblock regions. For each predicted haploblock, the local PCA MDS plot for the relevant chromosome, a PCA of the selected region, observed heterozygosity for each haploblock genotype, and LD patterns for the relevant chromosome are shown. In the local PCA MDS plots, each dot represents a 100 SNP window, and windows within the haploblock region are highlighted. The x-axis values represent Mbp. For *H. petiolaris*, haploblocks were identified in the full or subspecies datasets; the local PCA and LD plots are from the dataset where the haploblock was identified, while PCA and heterozygosity plots use the full dataset. In PCA plots, samples are colored by inferred haploblock genotype. For LD plots, upper triangle = all individuals; lower triangle = only individuals homozygous for the more common haploblock allele. Colours represent the second highest R^2^ value in 0.5 Mbp windows. For most haploblock regions, high LD is driven by differences between haplotypes, so high LD is removed when only one haplotype is present. Box plots show the median, box edges represent the 25^th^ and 75^th^ percentiles, whiskers represent the maximum/minimum data points within 1.5x interquartile range outside box edges.

**Extended Data Fig. 5:**
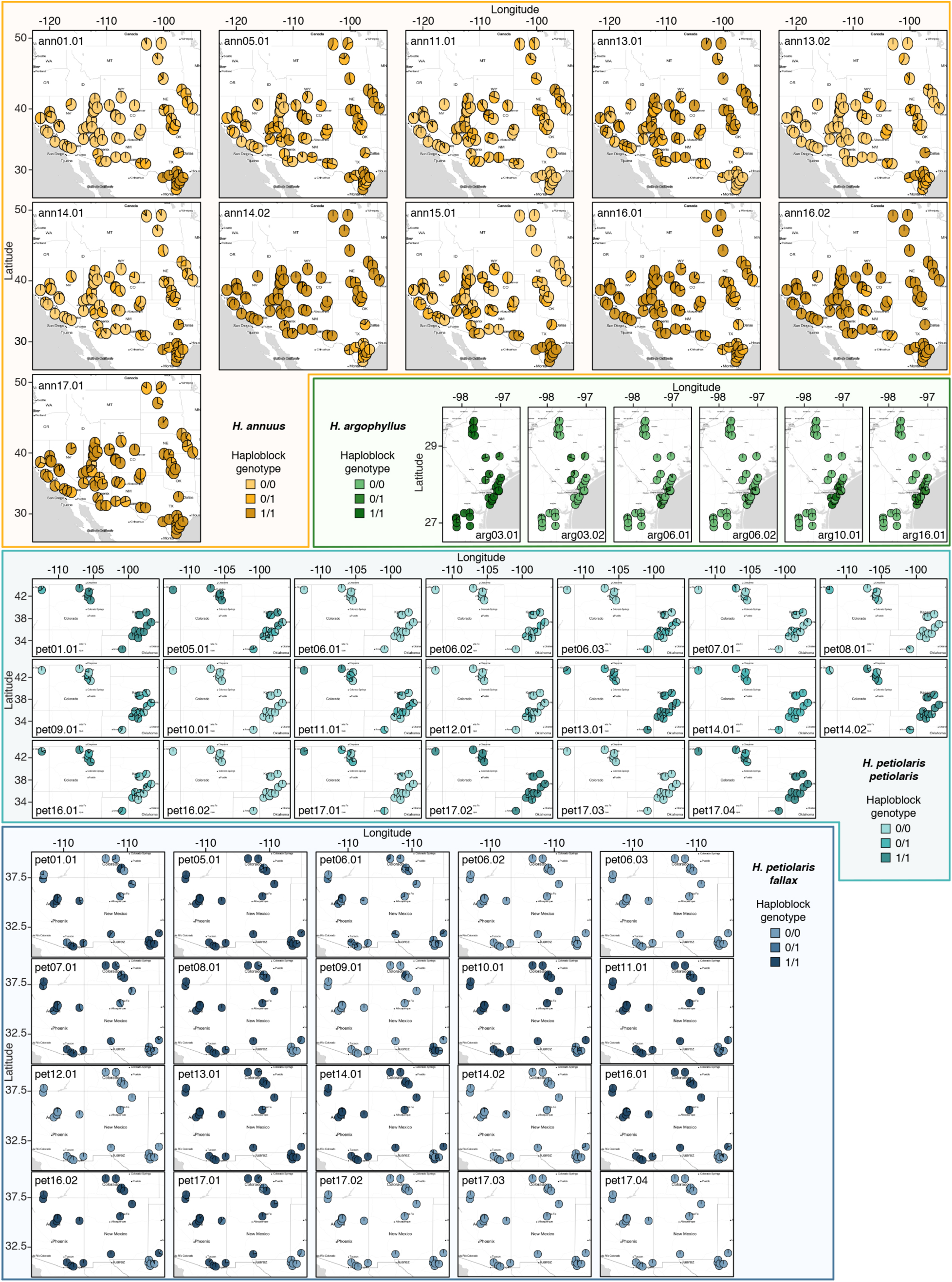
Geographic distribution of haploblock genotypes for the three sunflower species.

**Extended Data Fig. 6:**
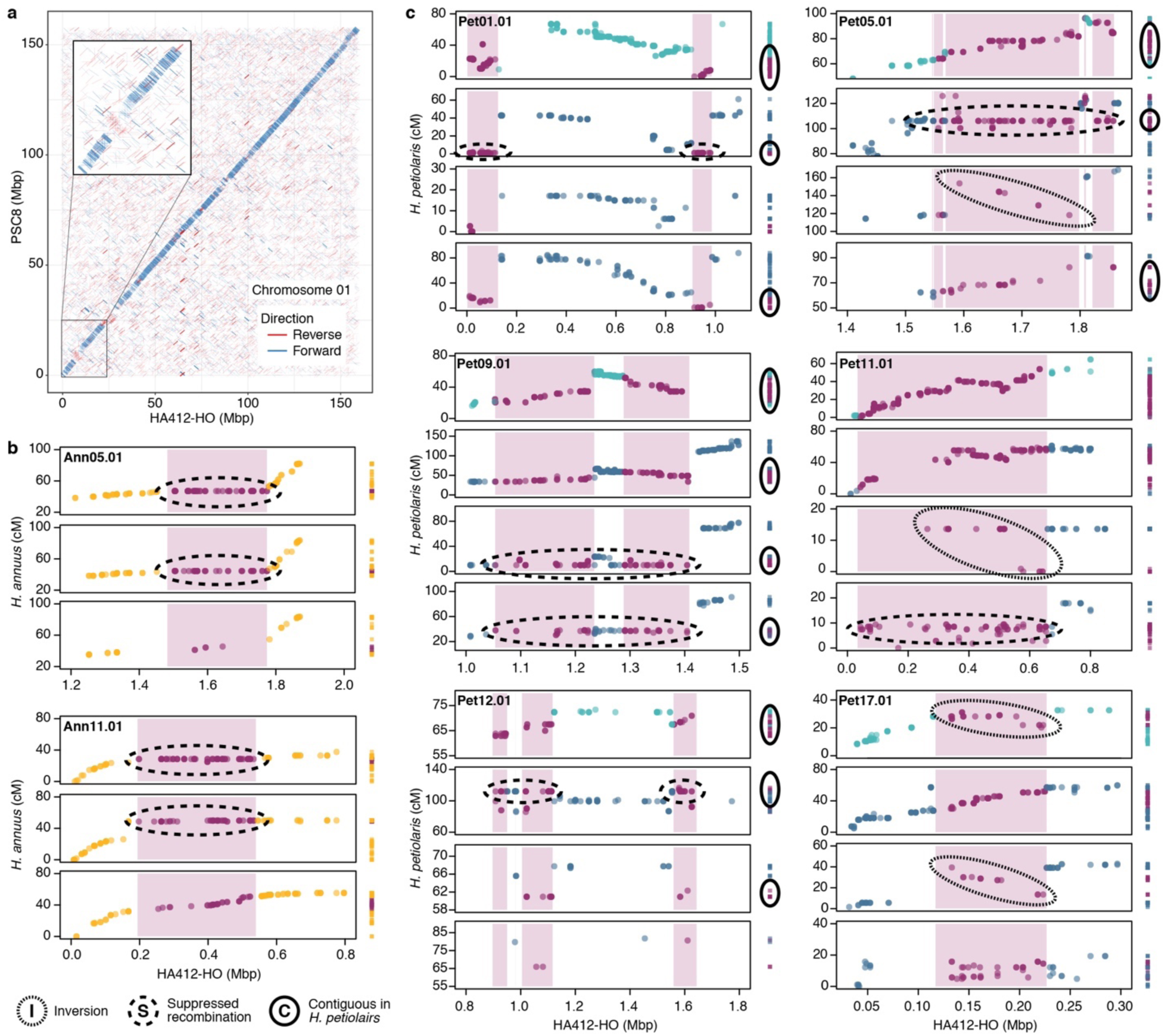
Comparisons between reference assemblies and genetic maps confirm structural rearrangements associated with haploblocks. **a**, Alignment of chromosome 1 for the *H. annuus* genome assemblies PSC8 and HA412-HOv2. The ann01.01 region (at ∼8 Mpb, see inset), for which the two cultivars have different haplotypes, shows inverted alignment. **b**, Three *H. annuus* genetic maps (constructed using F_2_populations between wild individuals and the HA412-HO cultivar) and **c**, Four genetic maps (constructed using F_1_ populations. From top to bottom: *H. petiolaris petiolaris*, *H. petiolaris fallax*^50^, newly constructed dune *H. petiolaris fallax*, and newly constructed non-dune *H. petiolaris fallax*) are plotted relative to the HA412-HOv2 reference assembly. To the right of each dot plot, markers are plotted in the order they appear in each genetic map. Haploblock regions and the markers that fall within them are highlighted in purple. Circled haploblock regions show evidence of different orientations across the multiple maps (dotted lines), of suppressed recombination (dashed lines), or are contiguous in *H. petiolaris* maps despite being split over multiple windows in the HA412-HOv2 reference assembly (solid lines). Parental haploblock genotypes are known for the *H. annuus* maps and for the bottom two *H. petiolaris* maps. Ann05.01 and ann11.01 were segregating within in the *H. annuus* mapping populations. Genotypes at pet05.01 and pet11.01 differed between the *H. petiolaris fallax* parents of newly constructed dune and non-dune populations, while both parents were heterozygous for the pet09.01 haploblock. In all these cases, patterns of segregation are consistent with the parental haploblock genotypes. For the remaining *H. petiolaris* maps, the parental haploblock genotypes are not known. Since an absence of evidence is uninformative in those cases, only haploblock regions with evidence for inversions or contiguous windows from these two maps are plotted.

**Extended Data Fig. 7:**
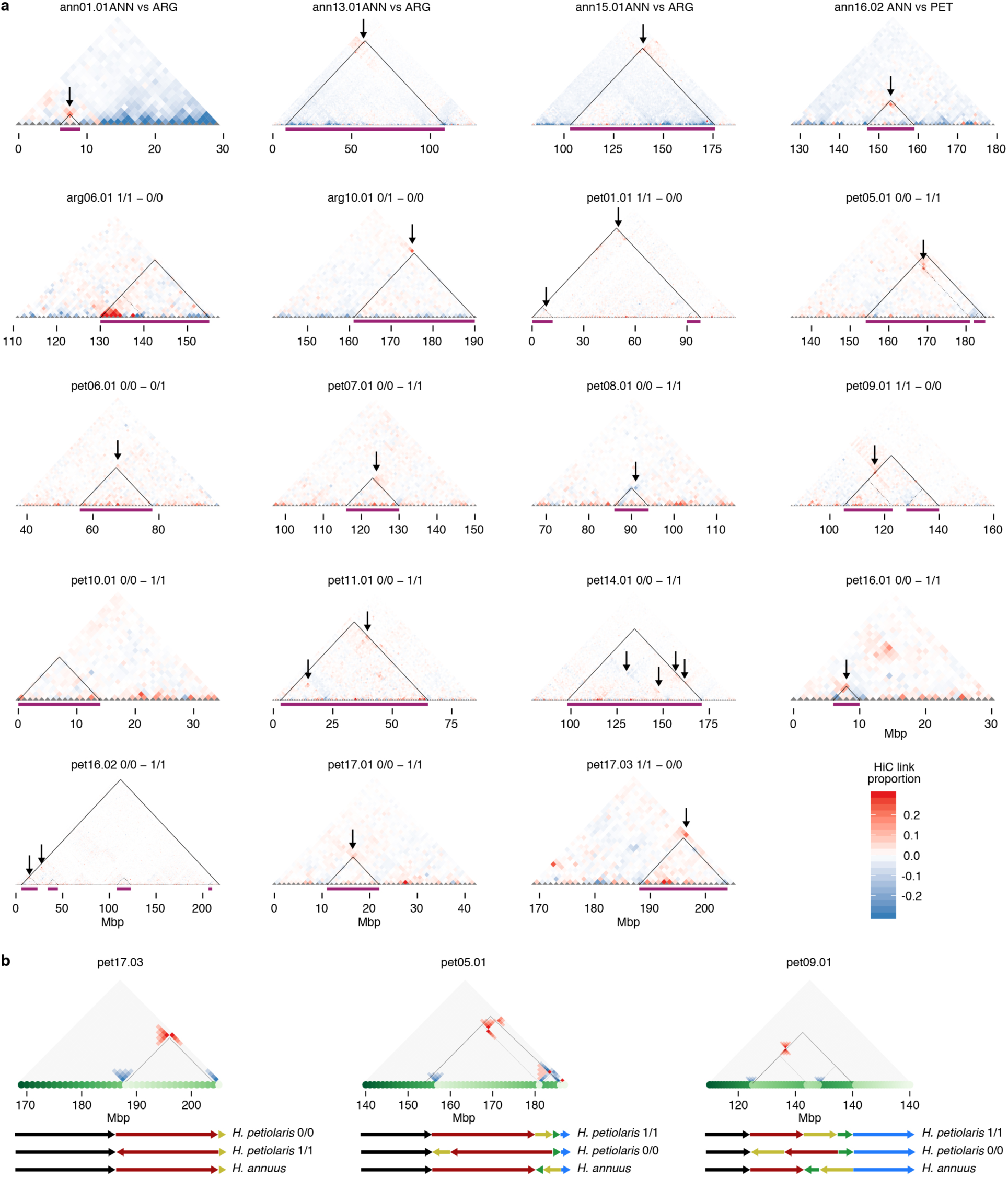
HiC comparisons identify SVs associated with most, but not all, haploblocks. **a**, Differences in HiC interactions between pairs of early- and late-flowering *H. argophyllus* or dune and non-dune *H. petiolaris* samples. Purple bars and solid black lines represent approximate haploblock boundaries. Pieces of a single haploblock that map to different regions of the HA412-HOv2 reference are highlighted by dotted lines. Top row = comparisons between *H. annuus* and *H. argophyllus* or *H. petiolaris*, for *H. annuus* haploblock regions. Since the relative haploblock genotypes between sunflower species are not known, only cases in which evidence of structural variance were observed are reported. Following rows = regions for which the pairs of *H. argophyllus* or *H. petiolairs* samples differed at haploblock alleles. Red or blue dots show increased long distance interactions in one sample, consistent with differences in genome structure. Relevant differences in long distance interactions are highlighted by black arrows. No evidence of large-scale structural variation was observed for arg06.01 and pet10.01. An excess of interactions in the early-flowering allele for the ∼130-140 Mbp region of chromosome 6 is consistent with the presence of deletions in the late flowering alleles (see Extended Data Fig. 2c), as well as with improved mappability of reads from the early flowering allele, which, being an introgression from wild *H. annuus*, is closer in sequence to the HA412-HO reference. Differences in HiC interactions were capped between -0.3 and 0.3 for plotting purposes. **b**, Inversion scenarios with comparisons of simulated HiC interaction matrixes consistent with empirical patterns. Note that there are *H. annuus*-specific inversions in the reference genome, as well as inversions between haploblocks.

**Extended Data Fig. 8:**
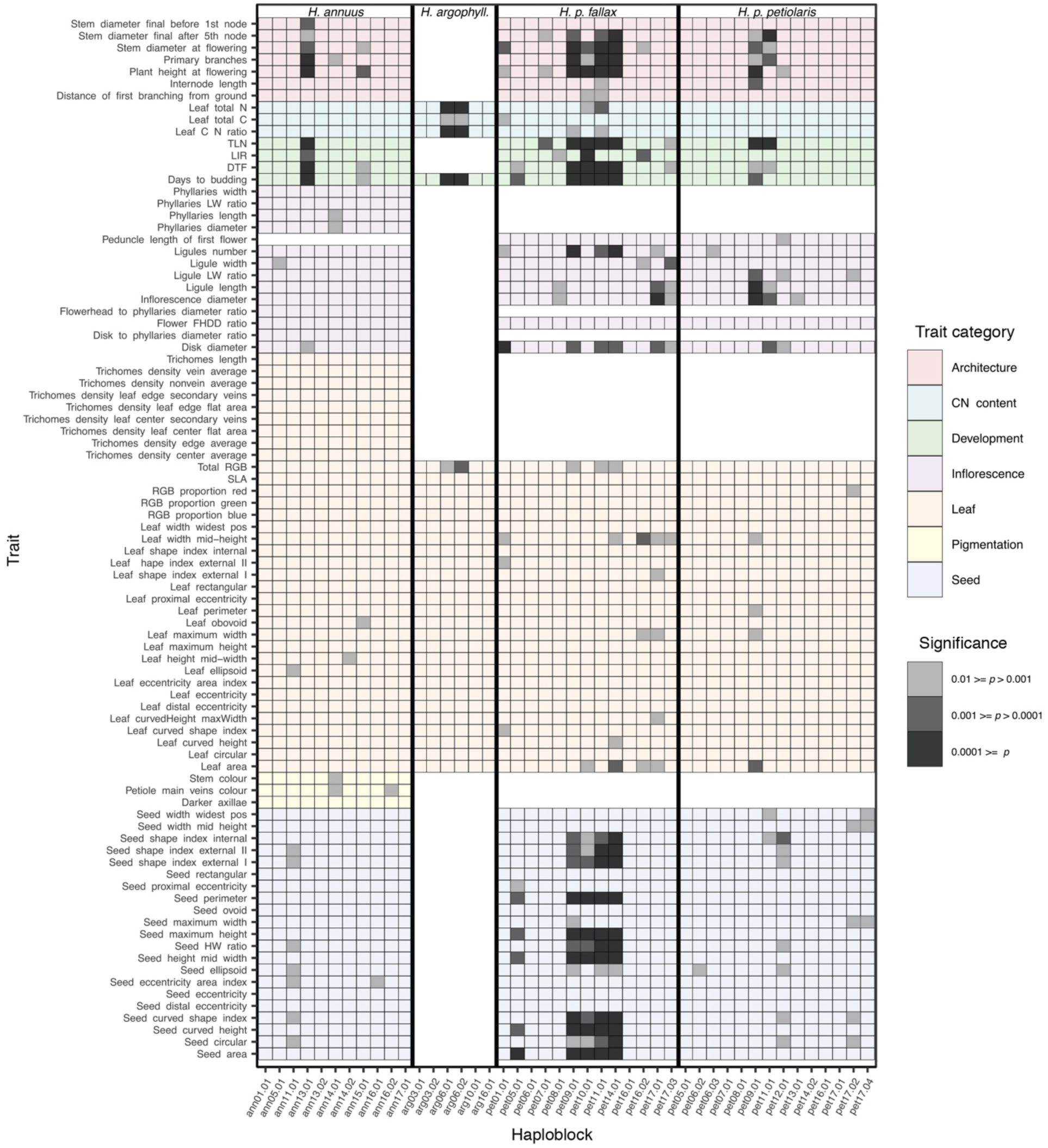
Haploblock GWAs. Heatmap of GWAs for individual phenotypic traits, treating haploblocks as individual loci. Haploblocks were filtered to retain only regions with minor allele frequency ≥ 3%. PCA and kinship matrices used as covariates were calculated without variants inside haploblock regions.

**Extended Data Fig. 9:**
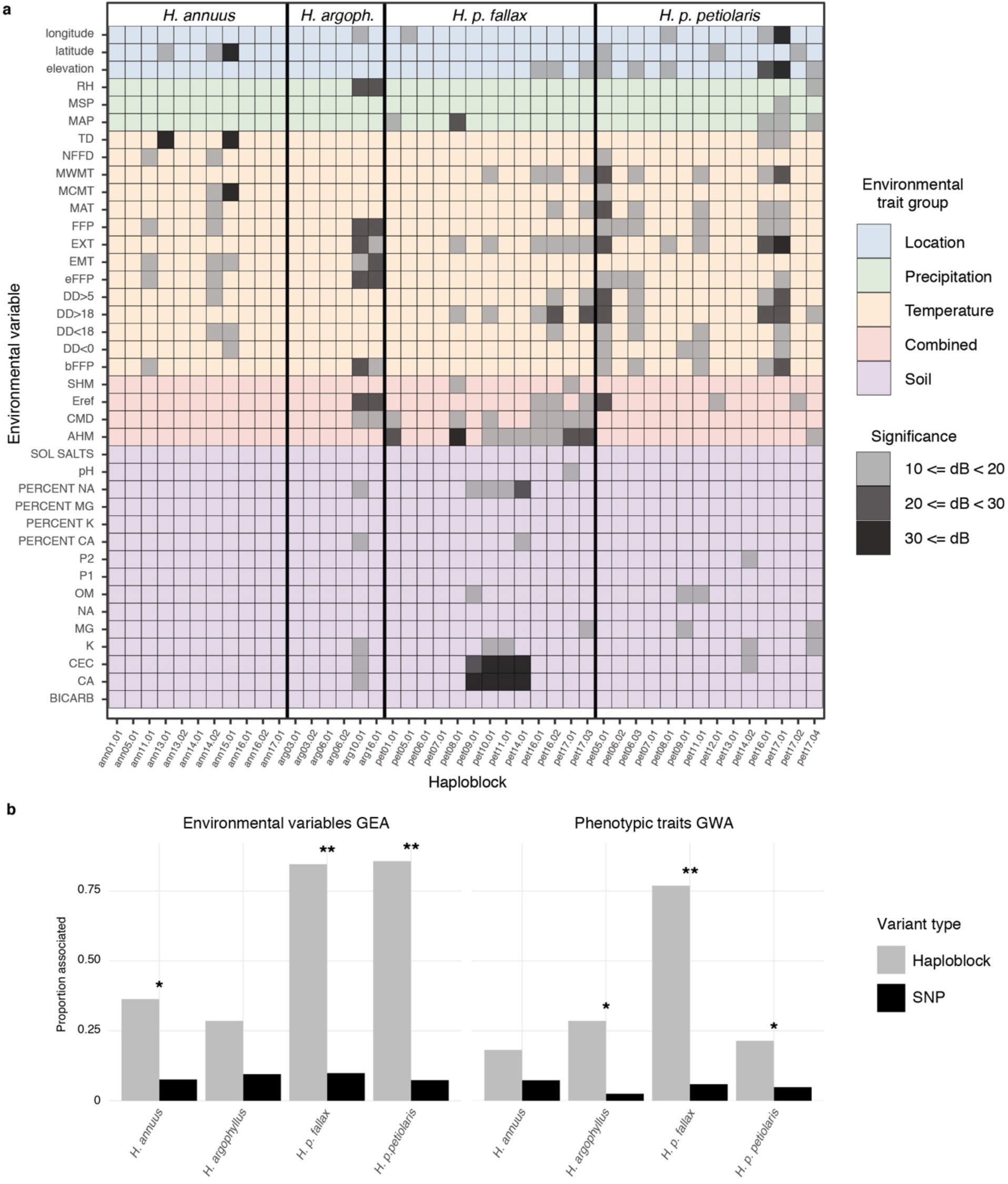
Haploblock GEAs. **a**, Heatmap of GEAs for individual environmental variables, treating haploblocks as individual loci. Haploblocks were filtered to retain only regions with minor allele frequency ≥ 3%. The population correlation matrix was calculated without variants inside haploblock regions. **b**, The proportion of haploblock and SNP loci significantly associated with one or more environmental variable (dB ≥ 10) or phenotypic trait (p ≤ 0.001). * p < 0.05, ** p < 0.0005 (two-sided proportion test).

## References

1 Romanes, G. J. Physiological selection; an additional suggestion on the origin of species. Zool. J. Linn. Soc. 19, 337–411 (1886).

2 Endler, J. A. Gene flow and population differentiation: studies of clines suggest that differentiation along environmental gradients may be independent of gene flow. Science 179, 243–250 (1973).

3 Felsenstein, J. Skepticism towards Santa Rosalia, or why are there so few kinds of animals? Evolution 35, 124–138 (1981).

4 Harter, A. V. et al. Origin of extant domesticated sunflowers in eastern North America. Nature 430, 201–205 (2004).

5 Whitney, K. D., Randell, R. A. & Rieseberg, L. H. Adaptive introgression of abiotic tolerance traits in the sunflower *Helianthus annuus*. New Phytol. 187, 230–239 (2010).

6 Ostevik, K. L., Andrew, R. L., Otto, S. P. & Rieseberg, L. H. Multiple reproductive barriers separate recently diverged sunflower ecotypes. Evolution 70, 2322–2335 (2016).

7 Moyers, B. T. The landscape of divergence in silverleaf sunflowers. PhD dissertation (2015).

8 Qiu, F. et al. Phylogenetic trends and environmental correlates of nuclear genome size variation in *Helianthus* sunflowers. New Phytol. 221, 1609–1618 (2019).

9 Badouin, H. et al. The sunflower genome provides insights into oil metabolism, flowering and Asterid evolution. Nature 546, 148–152 (2017).

10 Shagina, I. et al. Normalization of genomic DNA using duplex-specific nuclease. Biotechniques 48, 455–459 (2010).

11 Sunflower Genome Database, https://www.sunflowergenome.org/ (2019).

12 INRA Sunflower Bioinformatics Resources, https://www.heliagene.org/ (2019).

13 Baute, G. J., Owens, G. L., Bock, D. G. & Rieseberg, L. H. Genome-wide genotyping-by-sequencing data provide a high-resolution view of wild *Helianthus* diversity, genetic structure, and interspecies gene flow. Am. J. Bot. 103, 2170–2177 (2016).

14 Stephens, J. D., Rogers, W. L., Mason, C. M., Donovan, L. A. & Malmberg, R. L. Species tree estimation of diploid *Helianthus* (Asteraceae) using target enrichment. Am. J. Bot. 102, 910–920 (2015).

15 Heiser, C. B. & Smith, D. M. The North American sunflowers Helianthus. Published for the Club by the Seeman Printery, Durham, N.C., USA (1969).

16 Raduski, A. R., Rieseberg, L. H. & Strasburg, J. L. Effective population size, gene flow, and species status in a narrow endemic sunflower, *Helianthus neglectus*, compared to its widespread sister species, H. petiolaris. Int. J. Mol. Sci. 11, 492–506 (2010).

17 Strasburg, J. L. & Rieseberg, L. H. Molecular demographic history of the annual sunflowers *Helianthus annuus* and *H. petiolaris*—large effective population sizes and rates of long-term gene flow. Evolution 62, 1936–1950 (2008).

18 Blackman, B. K., Michaels, S. D. & Rieseberg, L. H. Connecting the sun to flowering in sunflower adaptation. Mol. Ecol. 20, 3503–3512 (2011).

19 Zan, Y. & Carlborg, Ö. A polygenic genetic architecture of flowering time in the worldwide *Arabidopsis thaliana* population. Mol. Biol. Evol. 36, 141–154 (2018).

20 Kobayashi, Y., Kaya, H., Goto, K., Iwabuchi, M. & Araki, T. A pair of related genes with antagonistic roles in mediating flowering signals. Science 286, 1960–1962 (1999).

21 Werner, J. D. et al. Quantitative trait locus mapping and DNA array hybridization identify an *FLM* deletion as a cause for natural flowering-time variation. Proc. Natl. Acad. Sci. USA 102, 2460–2465 (2005).

22 Cao, Y., Wen, L., Wang, Z. & Ma, L. SKIP interacts with the Paf1 complex to regulate flowering via the activation of *FLC* transcription in *Arabidopsis*. Mol. Plant 8, 1816–1819 (2015).

23 Wang, L. C. et al. Involvement of the *Arabidopsis* HIT1/AtVPS53 tethering protein homologue in the acclimation of the plasma membrane to heat stress. J .Exp. Bot. 62, 3609–3620 (2011).

24 Blackman, B. K. et al. Contributions of flowering time genes to sunflower domestication and improvement. Genetics 187, 271–287 (2011).

25 Brouillette, L. C. & Donovan, L. A. Nitrogen stress response of a hybrid species: a gene expression study. Ann. Bot. 107, 101–108 (2011).

26 Ostevik, K. L. The ecology and genetics of adpatation and speciation in dune sunflowers. PhD dissertation (2016).

27 Andrew, R. L. & Rieseberg, L. H. Divergence is focused on few genomic regions early in speciation: incipient speciation of sunflower ecotypes. Evolution 67, 2468–2482 (2013).

28 Li, H. & Ralph, P. Local PCA shows how the effect of population structure differs along the genome. Genetics 211, 289–304 (2019).

29 Ortiz-Barrientos, D., Engelstadter, J. & Rieseberg, L. H. Recombination Rate Evolution and the Origin of Species. Trends Ecol. Evol. 31, 226–236 (2016).

30 Trickett, A. J. & Butlin, R. K. Recombination suppressors and the evolution of new species. Heredity 73, 339 (1994).

31 Arostegui, M. C., Quinn, T. P., Seeb, L. W., Seeb, J. E. & McKinney, G. J. Retention of a chromosomal inversion from an anadromous ancestor provides the genetic basis for alternative freshwater ecotypes in rainbow trout. Mol. Ecol. 28, 1412–1427 (2019).

32 Joron, M. et al. Chromosomal rearrangements maintain a polymorphic supergene controlling butterfly mimicry. Nature 477, 203 (2011).

33 Lowry, D. B. & Willis, J. H. A widespread chromosomal inversion polymorphism contributes to a major life-history transition, local adaptation, and reproductive isolation. PLoS Biol. 8, e1000500, doi:10.1371/journal.pbio.1000500 (2010).

34 Fang, Z. et al. Megabase-scale inversion polymorphism in the wild ancestor of maize. Genetics 191, 883–894 (2012).

35 Wellenreuther, M., Rosenquist, H., Jaksons, P. & Larson, K. W. Local adaptation along an environmental cline in a species with an inversion polymorphism. J. Evol. Biol. 30, 1068–1077 (2017).

36 Belton, J. M. et al. Hi-C: a comprehensive technique to capture the conformation of genomes. Methods 58, 268–276 (2012).

37 Mason, C. M. How old are sunflowers? A molecular clock analysis of key divergences in the origin and diversification of *Helianthus* (Asteraceae). Int. J. Plant Sci. 179, 182–191 (2018).

38 Wellenreuther, M. & Bernatchez, L. Eco-evolutionary genomics of chromosomal inversions. Trends Ecol. Evol. 33, 427–440 (2018).

39 Kirkpatrick, M. & Barton, N. Chromosome inversions, local adaptation and speciation. Genetics 173, 419–434 (2006).

40 Lotterhos, K. E. The effect of neutral recombination variation on genome scans for selection. G3 (Bethesda) 9, 1851-1867 (2019).

41 Heiser Jr, C. B. Hybridization in the annual sunflowers: Helianthus annuus × H. debilis var. cucumerifolius. Evolution, 42–51 (1951).

42 Hooper, D. M. & Price, T. D. Chromosomal inversion differences correlate with range overlap in passerine birds. Nature Ecol. Evol. 1, 1526 (2017).

43 Clausen, J. Stages in the evolution of plant species. Cornell University Press, Ithaca, N.Y., USA (1951).

44 Heiser, C. B. Three new annual sunflowers (*Helianthus*) from the Southwestern United States. Rhodora 60, 272–283 (1958).

45 Andrew, R. L., Kane, N. C., Baute, G. J., Grassa, C. J. & Rieseberg, L. H. Recent nonhybrid origin of sunflower ecotypes in a novel habitat. Mol. Ecol. 22, 799–813 (2013).

46 Kirkpatrick, M. Reinforcement and divergence under assortative mating. Proc. Royal Soc. B 267, 1649–1655 (2000).

47 Feder, J. L., Gejji, R., Powell, T. H. & Nosil, P. Adaptive chromosomal divergence driven by mixed geographic mode of evolution. Evolution 65, 2157–2170 (2011).

48 Via, S. & West, J. The genetic mosaic suggests a new role for hitchhiking in ecological speciation. Mol. Ecol. 17, 4334–4345 (2008).

49 Yeaman, S. & Whitlock, M. C. The genetic architecture of adaptation under migration-selection balance. Evolution 65, 1897–1911 (2011).

50 Ostevik, K. L., Samuk, K. & Rieseberg, L. H. Ancestral reconstruction of sunflower karyotypes reveals dramatic chromosomal evolution. BioRxiv, doi:10.1101/737155 (2019).

## Methods references

51 Rowan, B. A., Patel, V., Weigel, D. & Schneeberger, K. Rapid and inexpensive whole-genome genotyping-by-sequencing for crossover localization and fine-scale genetic mapping. G3 (Bethesda) 5, 385–398 (2015).

52 Rohland, N. & Reich, D. Cost-effective, high-throughput DNA sequencing libraries for multiplexed target capture. Genome Res. 22, 939–946 (2012).

53 Matvienko, M. et al. Consequences of normalizing transcriptomic and genomic libraries of plant genomes using a duplex-specific nuclease and tetramethylammonium chloride. PLoS One 8, e55913, doi:10.1371/journal.pone.0055913 (2013).

54 Hubner, S. et al. Sunflower pan-genome analysis shows that hybridization altered gene content and disease resistance. Nat. Plants 5, 54–62 (2019).

55 Lee-Yaw, J. A., Grassa, C. J., Joly, S., Andrew, R. L. & Rieseberg, L. H. An evaluation of alternative explanations for widespread cytonuclear discordance in annual sunflowers (*Helianthus*). New Phytol. 221, 515–526 (2019).

56 Owens, G. L., Baute, G. J., Hubner, S. & Rieseberg, L. H. Genomic sequence and copy number evolution during hybrid crop development in sunflowers. Evol. Appl. 12, 54–65 (2019).

57 Bolger, A. M., Lohse, M. & Usadel, B. Trimmomatic: a flexible trimmer for Illumina sequence data. Bioinformatics 30, 2114–2120 (2014).

58 Sedlazeck, F. J., Rescheneder, P. & von Haeseler, A. NextGenMap: fast and accurate read mapping in highly polymorphic genomes. Bioinformatics 29, 2790–2791 (2013).

59 Poplin, R. et al. Scaling accurate genetic variant discovery to tens of thousands of samples. BioRxiv, doi:10.1101/201178 (2017).

60 Li, H. Aligning sequence reads, clone sequences and assembly contigs with BWA-MEM. ArXiv, arXiv:1303.3997v1302 (2013).

61 Danecek, P. et al. The variant call format and VCFtools. Bioinformatics 27, 2156–2158 (2011).

62 Hoang, D. T., Chernomor, O., von Haeseler, A., Minh, B. Q. & Vinh, L. S. UFBoot2: Improving the Ultrafast Bootstrap Approximation. Mol. Biol. Evol. 35, 518–522 (2018).

63 Kalyaanamoorthy, S., Minh, B. Q., Wong, T. K. F., von Haeseler, A. & Jermiin, L. S. ModelFinder: fast model selection for accurate phylogenetic estimates. Nat. Methods 14, 587–589 (2017).

64 Nguyen, L. T., Schmidt, H. A., von Haeseler, A. & Minh, B. Q. IQ-TREE: a fast and effective stochastic algorithm for estimating maximum-likelihood phylogenies. Mol. Biol. Evol. 32, 268–274 (2015).

65 Grimm, D. G. et al. easyGWAS: A Cloud-Based Platform for Comparing the Results of Genome-Wide Association Studies. Plant Cell 29, 5–19 (2017).

66 Wang, T., Hamann, A., Spittlehouse, D. & Carroll, C. Locally Downscaled and Spatially Customizable Climate Data for Historical and Future Periods for North America. PLoS One 11, e0156720, doi:10.1371/journal.pone.0156720 (2016).

67 Gautier, M. Genome-Wide Scan for Adaptive Divergence and Association with Population-Specific Covariates. Genetics 201, 1555–1579 (2015).

68 Jeffreys, H. Theory of Probability. Oxford University Press, London/New York/Oxford (1961).

69 Hellens, R. P., Edwards, E. A., Leyland, N. R., Bean, S. & Mullineaux, P. M. pGreen: a versatile and flexible binary Ti vector for Agrobacterium-mediated plant transformation. Plant Mol. Biol. 42, 819–832 (2000).

70 Weigel, D. & Glazebrook, J. Arabidopsis: A laboratory manual. CSHL Press, Cold Spring Harbor, N.Y., USA (2002).

71 Kurtz, S. et al. Versatile and open software for comparing large genomes. Genome Biol. 5, R12 (2004).

72 Goel, M., Sun, H., Jiao, W.-B. & Schneeberger, K. SyRI: finding genomic rearrangements and local sequence differences from whole-genome assemblies. BioRxiv, doi:10.1101/546622 (2019).

73 Marie-Nelly, H. et al. High-quality genome (re)assembly using chromosomal contact data. Nat. Commun. 5, 5695 (2014).

74 Heinz, S. et al. Simple combinations of lineage-determining transcription factors prime cis-regulatory elements required for macrophage and B cell identities. Mol. Cell 38, 576–589 (2010).

75 R Core Team. R: A language and environment for statistical computing, https://www.R-project.org/ (2019).

76 Suchard, M. A. et al. Bayesian phylogenetic and phylodynamic data integration using BEAST 1.10. Virus Evol. 4, vey016, doi:10.1093/ve/vey016 (2018).

77 Rambaut, A., Drummond, A. J., Xie, D., Baele, G. & Suchard, M. A. Posterior summarization in Bayesian phylogenetics using Tracer 1.7. Syst. Biol. 67, 901–904 (2018).

78 Sambatti, J. B., Strasburg, J. L., Ortiz-Barrientos, D., Baack, E. J. & Rieseberg, L. H. Reconciling extremely strong barriers with high levels of gene exchange in annual sunflowers. Evolution 66, 1459–1473 (2012).

79 Yu, G., Smith, D. K., Zhu, H., Guan, Y. & Lam, T. T. Y. ggtree: an R package for visualization and annotation of phylogenetic trees with their covariates and other associated data. Methods Ecol. Evol. 8, 28–36 (2017).

80 Rambaut, A. FigTree, http://tree.bio.ed.ac.uk/software/figtree/ (2009).

