## Supplementary Methods for "Massive haplotypes underlie ecotypic differentiation in sunflowers"

### Common garden

Ten mother plants were randomly selected from each of the 151 populations that were included in the common garden experiment. Ten seeds from each of these plants were surface-sterilized by immersing them for 10 minutes in a 1.5% sodium hypochlorite solution. Seeds were then rinsed twice in distilled water and treated for at least one hour in a solution of 1% PPM (Plant Cell Technologies, Washington, DC, USA), a broad-spectrum biocide/fungicide, to minimize contamination, and 0.05 mM gibberellic acid (Sigma-Aldrich, St. Louis, MO, USA). They were then scarified, de-hulled, and kept for two weeks at 4 ºC in the dark on filter paper imbibed with a 1% PPM solution. Following this, seeds were kept in the dark at room temperature until they germinated, and then transplanted in peat pots. Seedlings were grown in a greenhouse for two weeks and then moved to an open-sided greenhouse for a week for acclimation. Plants were transplanted into three separate fields (one for each sunflower species) at the Totem Plant Science Field Station of the University of British Columbia (Vancouver, Canada) on the 25$^{th}$ of May (*H. argophyllus*), 2$^{nd}$ of June (*H. petiolaris*) and 7$^{th}$ of June 2016 (*H. annuus*). Within each field, pairs of plants from the same population of origin were sown using a completely randomized design. Phenotypic data were collected throughout the growing season, as detailed in Supplementary Table 1. Morphometric data were extracted from digital pictures using Fiji[1,2] and Tomato Analyzer[3]. Plants were grown until the beginning of November, by which point almost all the plants had flowered.

**DNA isolation, library preparation and sequencing**

Tissue from young leaves was collected from all individual plants, and genomic DNA was extracted from leaf tissue using a CTAB protocol (modified from Murray and Thompson, 1980[4] and Zeng *et al.* 2002[5]), the DNeasy Plant Mini Kit or a DNeasy 96 Plant Kit (Qiagen, Hilden, Germany). DNA was sheared to an average fragment size of 400 bp using a Covaris M220 ultrasonicator (Covaris, Woburn, Massachusetts, USA), following the manufacturer's recommendations. 750 ng of sheared DNA were used as starting material to prepare paired-end whole-genome shotgun (WGS) Illumina libraries for 719 *H. annuus*, 488 *H. petiolaris*, 299 *H. argophyllus* individuals, and twelve additional samples from annual and perennial sunflowers (Supplementary Table 1), using a protocol largely based on Rowan *et al*, 2015[6], the TruSeq DNA Sample Preparation Guide from Illumina (Illumina, San Diego, CA, USA) and Rolhand *et al.* 2012[7]. End-repairing of the sheared DNA fragments was performed using the NEBNext End Repair Module (NEB, Ipswich, Massachusetts, USA). The fragments were then A-tailed using Klenow Fragment (3'-->5'exo-; NEB) and ligated to 24-bp-long, non-barcoded adapters with a 3' T-overhang using the Quick Ligation Kit (NEB). After each enzymatic step, the reactions were purified using 1.6 volumes of paramagnetic SPRI beads, prepared according to Rohland *et al.* 2012[7]. An enrichment step was then performed using KAPA HiFi HotStart ReadyMix

(Roche, Basel, Switzerland) and short, non-indexed primers that do not extend the adapters. The reactions were then purified using 1.6 volumes of SPRI beads.

The sunflower genome contains a very large amount of highly repetitive sequences derived from the recent expansion of two retrotransposon families[8]. In order to reduce the representation of repetitive sequences, including plastid DNA, the enriched libraries were treated with a Duplex-Specific Nuclease (DSN; Evrogen, Moscow, Russia), following the protocols reported in Shagina *et al.* 2010[9] and Matvienko *et al.* 2013[10], with modifications. Depletion conditions were optimized for the sunflower genome by quantitative PCR; relative abundance of chloroplast DNA and transposable elements before and after depletion was estimated using a primer pair recognizing a chloroplast gene, and degenerate primers recognizing one of the most abundant transposon families in the sunflower genome, and comparing them to the abundance of the single copy *HaLFY* gene. Libraries were concentrated using SPRI beads to a concentration of 160 ng/µl. Three µl of libraries were mixed to 1 µl of hybridization buffer (200 mM HEPES pH 7.5, 2 M NaCl, 0.8 mM EDTA), overlaid with 10 µl of mineral oil, and incubated at 78 ºC for 22 hours. Five µl of pre-warmed DSN buffer (0.1 M Tris pH 8.0, 10 mM $MgCl_2$, 2 mM DTT) were then added to each sample. After a five minutes incubation at 70 ºC, 0.1 U of DSN enzyme was added to the samples, and they were incubated for a further 15 minutes at 70 ºC. Digestion was stopped by adding 10 µl of 10 mM EDTA. The fragments were then further amplified using KAPA HiFi HotStart ReadyMix (Roche, Basel, Switzerland) and primers that completed the adapters and added a six-base pair index to the P7 adapter. All adapter and primer sequences are reported in Supplementary Table 3.

After amplification, the libraries were purified with 1 volume of SPRI beads, quantified using a QuBit dsDNA Broad Range Assay Kit (Invitrogen, Carlsbad, California, USA) and analyzed on a 2100 Bioanalyzer instrument using a High Sensitivity DNA Analysis Kit (Agilent, Santa Clara, California, USA).

**Variant calling**

The call set comprised a total of 2392 samples (Supplementary Table 1). Illumina adapters and poor quality reads were hard-clipped using Trimmomatic[11] (v0.36). Reads shorter than 36 bp at this step were dropped. All remaining reads (including orphaned reads with a pair) were then aligned to the *H. annuus* XRQv1[12] genome (HanXRQr1.0-20151230) using NextGenMap[13] (v0.5.3). The aligner produced three mapped sam files (mapped pairs, mapped unpaired forward reads, and mapped unpaired reverse reads), which were converted to BAM, concatenated, and then sorted (samtools[14,15] v0.1.19). To finalize the alignment, PCR duplicates were marked (picard[16] MarkDuplicates 2.9.3) and the BAM file was indexed. Some sequencing libraries were sequenced in multiple lanes to increase coverage; BAM files for a same individual were merged by sample ID (sambamba[17] v0.6.6) and PCR duplicates were remarked.

We implemented the Genome Analysis ToolKit (GATK 4.0.1.2) germline short variant discovery pipeline to perform variant calling[18]. To reduce computational time and improve variant quality, we excluded genomic regions containing transposable elements, which represent ~¾ of the sunflower genome, and to which short reads cannot be reliably mapped. These callable regions comprised 1.1 GB of the total 3.6 Gbp of the XRQv1 assembly[12]; the corresponding bed file is included in the code repository (HanXRQr1.0-20151230_allTEs_ubc.non-repetitive-regions.2017.sorted.bed). All downstream analyses were conducted on this TE-filtered dataset. HaplotypeCaller (v4.0.1.2) was used on each sample individually to produce a Genomic VCF (g.vcf). Heterozygosity settings for HaplotypeCaller step were increased to mu = 0.01 and st_dev

= 0.1. This is 10 fold higher than the default, but better reflects the expected diversity in sunflowers compared to humans.

HaplotypeCaller is a compute-intensive process that can take advantage of parallelism. To speed up the HaplotypeCaller phase, the callable regions of the genome were evenly split into 160 contiguous, non-overlapping genomic intervals. For each sample, those intervals were then processed in parallel, according to the number of cores available on the compute node. The 160 resulting genomic VCFs (.g.vcf) were then gathered into a single per-sample g.vcf, and then indexed using tabix and bgzip (v0.2.5-0). Joint genotyping of all samples in the same VCF would be ideal, as it allows for greater confidence on low frequency variants and simplifies comparisons between groups of samples. An initial attempt to jointly genotype all samples for 10 random 1-Mbp windows completed; however, given the large number of samples, high levels of genetic variation, and large genome size, it would have been computationally difficult to carry this operation across the genome given the available resources. Samples were therefore subdivided by species in three cohorts: *H. annuus*, *H. argophyllus* and *H. petiolaris*. Each cohort was independently genotyped.

Before further analysis, the g.vcf files were converted into a modified TileDB format[19] using GATK's GenomicsDBImport (v4.0.1.2). This step aggregates variants in a genomic region of interest from all samples in a cohort, and was found to be necessary to allow the next steps in the analysis to proceed. This operation was parallelized over 4 Mbp regions of the genome. TileDBs for a given region across a cohort were then converted into an unfiltered VCF using GATK's GenotypeGVCFs (v4.0.1.2) in mode `--use-new-qual`. The new-qual mode is the default mode in newer versions of GATK (≥4.1.1.0), and was necessary to allow SNP calling to run on our compute nodes (32- or 48-core Intel Skylake, with ≤256GB of RAM). Raw VCF chunks were then gathered into roughly per-chromosome files (17 files, one for each nuclear chromosome, plus one bundle file for all "unplaced" chromosome contigs HanXRQChr00c*, chloroplast, and mitochondria) using GatherVcfs (v4.0.1.2).

**Variant quality filtering**

The resulting raw VCF files contained an extremely large number of variant sites (222, 78 and 167 million variants for *H. annuus*, *H. argophyllus* and *H. petiolaris* respectively, combining SNPs and indels). The proportion of multi-allelic variant sites was strikingly high, varying between 24% and 51% across cohorts. We used GATK's recommended VariantRecalibrator to remove low-quality calls and produce a dataset of a more manageable size. VariantRecalibrator uses one or more "truth sets" of externally validated variant sites to decide which variants from a call set are likely to be real or artifacts. The model computed by the recalibrator attempts to define boundaries in the multidimensional site quality space that capture all or most known variant sites. Unknown variants that fall within this boundary are included, while those outside of the boundary are removed. In this way, stringency is determined by choosing the proportion of the known sites to be included in the boundary, which in GATK nomenclature is called the tranche. By selecting a smaller tranche (e.g. choosing tranche 90% over tranche 99%), the model selects a more stringent boundary and produces a smaller number of more confident sites. As a measure of variant quality, GATK measures the transition-transversion ratio for each tranche; this value was found to be ~2.1 in humans, although it is known to vary between species and between genomic regions[20,21] (Note: GATK uses 2.15 as a default target value for this metric).

While widely validated truth sets are available for humans, no such set exists to date for sunflower. We therefore defined a "gold set" using variants from samples in our dataset with the highest sequencing coverage. For *H. annuus*, this sample set included the top 67 inbred cultivar lines from the SAM population[22], and for *H. petiolaris* and *H. argophyllus* it included the top 20 wild samples. Variants were then filtered using the following parameters: Mapping Quality > 50.0, 90% sample coverage for the site, -1.0 > Strand odds Ratio < 1.0, Minor Allele Frequency > 0.25, Excess heterozygosity < 5.0 (for non-cultivar lines < 10.0 was used), -1.0 > BaseQRankSum < 1.0, Depth of coverage within one standard deviation from the mean and Excess Het > -4.5. The gold set was then recalibrated against the set of all variants from the entire corresponding cohort, using VariantRecalibrator (v4.0.6.0, with resource parameters `known = false, training = true, truth = true, prior = 10.0`). To speed up processing time, and to bring memory requirements to practical levels (i.e. < 250GB), it was necessary to pre-process the large training set before calibration; we stripped genotype information columns (with MakeSitesOnlyVcf) since the genotype columns from the VCF are not consulted by VariantRecalibrator. Following recommended practices, an early filtering pass to remove sites with extremely unlikely heterozygosity (ExcessHet z-score < 4.5) was also performed.

The 90% tranche for each cohort was selected for further analyses, based on the trade-off between SNP number and improvement to transition/transversion ratio. The full raw set of variants for each cohort was hard filtered according to this 90% tranche, and all indels were removed. Filtering by tranche retained 13.1%, 24.5%, and 30.7% of the total raw snps for *H. annuus, H. petiolaris, and H. argophyllus,* respectively.

After filtering for variant quality (i.e. 90% tranche) in each species, we subset samples into smaller sets for individual analyses and applied an additional filter. For each subset, variants were filtered to retain only bi-allelic SNPs with minor allele frequency ≥ 1% and genotype rate ≥

90%. The samples included in subsets used for different analyses (GWA, GEA) are listed in Supplementary Table 1.

The pipeline described in this section, including its data and software dependencies, were programmed into a Snakemake[23] (v4.7.0) workflow. To ensure reproducibility, the pipeline also makes extensive use of conda package environments, and Docker containers with precise versioning. Calling and filtering was computed on Compute-Canada's High-Performance-Computing (HPC) Cedar cluster.

**Remapping sites to the HA412-HO reference genome**

To remap our variant locations, 200 bp of reference sequence flanking each site in XRQv1 were extracted and aligned to HA412-HOv2 using BWA[24]. These alignments were filtered for mapping quality > 40 and the HA412-HOv2 position for the variant site was extracted. Since all remapped sites were not in repetitive regions and had passed VQSR filtering, remapping success rate was high (96-98%). Whenever mapping suggested two different variants on the XRQv1 genome were in the same position on the HA412-HOv2 genome, likely due indels and imprecise alignment, one site was shifted by one bp so they did not overlap. Remapping was preferred to *de novo* read alignment and variant calling against the HA412-HOv2 assembly because of the prohibitive amount of computational time that would have required. To test whether remapping improved the representation of linkage patterns across the genome, $R^2$ between all sites within 200kb on chromosome 2 was calculated using vcftools[25].

**Phylogenetic analysis**

Based on the results of the phylogenetic analysis, cases in which samples grouped outside their assumed population or species were reassigned if a source of error was confidently identified (i.e. mis-labelling during DNA extraction, library preparation or sequence analysis). Otherwise, the sample was removed. Note that samples with more intermediate phylogenetic positions were not removed, since they could represent admixed ancestry rather than mis-identification.

**Genome-wide association mapping**

Samples that were sequenced but were not part of the common garden experiment were removed from the variants dataset before filtering for minor allele frequency. Variants used for association were initially filtered for VQSR 90% tranche, and then further filtered to only include bi-allelic SNPs genotyped in ≥ 90% of samples and with a minor allele frequency ≥ 3%. While initially this dataset was mapped to the XRQv1 reference genome, all presented analyses use the remapped HA412-HOv2 genome positions. Variants were imputed and phased using Beagle[26] (version 10Jun18.811). Population structure was controlled for by including the first three principal components as covariates, as well as an IBS kinship matrix calculated by EMMAX[27]. We ran each trait GWA using EMMAX (v07Mar2010), as well as the EMMAX module in EasyGWAS[28]. EasyGWAS permits the use of different SNP dominance encoding and public release of all GWA results in an interactive format (https://easygwas.ethz.ch/gwas/results/xxx/), while the command line version of EMMAX allowed for faster batch submission and processing of results. Both approaches use the same method and produced comparable results. For every SNP/peak above the Bonferroni significance threshold, genes within a 100 kbp interval centered in the SNP with the lowest p-value, or within the boundaries of the GWA peak (whichever is larger), are reported in Supplementary Table 2.

**Genome-environment association analyses**

Population structure was estimated by choosing 10,000 putatively neutral random SNPs under the BayPass core model[29]. The Bayes factor (denoted $BF_{is}$ as in Gautier, 2015[29]) was then calculated under the standard covariate model to evaluate the association of SNP frequencies with 39 geographic, climatic and soil variables. For each SNP, $BF_{is}$ was expressed in deciban units [dB, $10\log_{10}$ ($BF_{is}$)]. Population PET_30 was removed from GEA analyses of *H. petiolaris petiolaris*, since very divergent haplotypes on two chromosomes made it an extreme outlier in the population correlation matrix, which resulted in GEA association values that were overall much lower than in the other three datasets. Populations ANN_71 and PET_21 were removed from the soil GEA analyses because no soil samples were available for them.

To calculate a significance threshold for candidate gene identification, pseudo-observed data (POD) were employed with the random 10,000 SNPs used for the core model, and a 1% empirical threshold was calculated for the observed Bayes factor. This value ranged from 6.7 to 7.3 depending on the species, and produced an extremely large number of outlier regions. We therefore followed Gautier, 2015[29] and employed Jeffreys' rule[30], quantifying the strength of associations between SNPs and variables as "strong" (10 dB $\leq BF_{is} <$ 15 dB), "very strong" (15 dB $\leq BF_{is} <$ 20 dB) and "decisive" ($BF_{is} \geq$ 20 dB). To produce a narrower set of candidate genes, the top ten non-overlapping 50 SNP windows based on the median $BF_{is}$ value were selected for each species and variable. A list of all the genes within these windows with at least one SNP with $BF_{is} \geq$ 20 dB within 1 kbp of their boundaries is reported in Supplementary Table 2.

**Transgenes and expression assays**

Total RNA was isolated from mature leaves and apical meristems using TRIzol (Thermo Fisher Scientific, Waltham, MA, USA) and cDNA was synthesized using the RevertAid First Strand cDNA Synthesis kit (Thermo Fisher Scientific).

**Population genomic detection of haploblocks**

Potential haploblock regions were defined based on MDS plots, and an MDS axis and minimum or maximum value that included windows within the region, but excluded the rest of the chromosome, were manually selected. Since there was variation in MDS score within each region, and an individual window within the region may fall below the cut off, windows that were surrounded by selected windows, within a range of 20 windows, were included. In most cases this resulted in a single unbroken range, but some regions, mainly *H. argophyllus* and *H. petiolaris*, were broken into multiple nearly abutting ranges. Furthermore, for *H. petiolaris* several of the regions were broken into unconnected distant regions, which likely reflects rearrangements in the *H. petiolaris* genome relative to the *H. annuus* reference used (see also Extended Data Fig. 6c).

All SNPs within the regions defined by MDS scores were used to calculate PCAs using SNPrelate[31]. The k-means clustering algorithm in R was used to define three clusters from PC1[32,33]. Since sample sizes were often unbalanced between the three potential groups, the starting positions for the three clusters were chosen as the maximum, minimum and middle of the range of PC1 scores. K-means cluster assignment was used as a preliminary genotype for the sample. Observed heterozygosity was also measured in each group. For all retained regions, samples clearly fell into three groups and observed heterozygosity was higher in the middle (0/1) group.

To visualize LD patterns, all SNPs with minor allele frequencies <5% were removed, the remaining variants were thinned to one per 100 bp, and genotype $R^2$ values for all sites within a chromosome were calculated. Values were grouped into 500 kb windows and the second largest $R^2$ value was plotted (Fig. 4e; Extended Data Fig. 4). In each case, regions identified in lostruct had high LD. The underlying recombination landscape in haploblock regions was explored by subsetting our dataset to samples homozygous for the more common haploblock genotype and measuring LD across the region. As before, SNPs with minor allele frequencies <5% were removed, variants were thinned to one per 100 bp, and genotype $R^2$ values for all sites within a chromosome were calculated. If the signal of high LD is only present when both haploblock genotypes are included, then it supports mechanisms that specifically prevent recombination between haplotypes. That being said, some haploblocks fall in generally low recombination regions and high LD within a haploblock genotype does not preclude recombination suppression.

**Synchronizing haploblocks in *H. petiolaris* subspecies.**

Lostruct was run in SNP datasets containing *H. petiolaris petiolaris, H. petiolaris fallax,* and both subspecies together. Although each dataset produced a collection of haploblocks, they were not identical. Some haploblocks were identified in one subspecies, but not the other, and some were only identified when both subspecies were analyzed together. In some cases, it was clear that haploblocks identified in both subspecies represented the same underlying haploblock because they physically overlapped and had overlapping diagnostic markers. We manually curated the list of haploblocks and merged those found in multiple datasets. We set the boundaries of these merged haploblocks to be inclusive (i.e. include windows found in either) and the diagnostic markers to be exclusive (i.e. only include sites found in both). For this merged set of haploblocks, all *H. petiolaris* samples were genotyped using diagnostic markers.

**Hi-C**

Based on our re-sequencing data, a pair of *H. petiolaris* and a pair *H. argophyllus* populations were selected that diverged for the largest number of haploblocks (PET_47 and PET_08 for *H. petiolaris* and ARG_18 and ARG_23 for *H. argophyllus*). Several individuals from each population were grown and genotyped at diagnostic SNPs for several haploblocks (pet09.01, pet10.01, pet10.01 and pet14.01 for *H. petiolaris*; arg06.01 and arg10.01 for *H. argophyllus*) using cleaved-amplified polymorphic sequence (CAPS) markers or direct Sanger sequencing (primers are reported in Supplementary Table 3). Chromosome conformation capture sequencing[34,35] (Hi-C) was then performed on one individual each from these four populations, to compare the structural organization of the different haplotypes at haploblock regions. Additionally, three Hi-C libraries sequenced on one HiSeq X lane from *H. annuus* HA412-HO were included in the analysis. This data was used to assemble the current HA412-HOv2 reference genome[36], and is used here as an interaction baseline.

Hi-C libraries were prepared by Dovetail Genomics (Scotts Valley, CA, USA) using the four-cutter restriction enzyme DpnII. Given the size and repetitive nature of sunflower genomes, Hi-C data could not be used to assemble a full genome; the HA412-HOv2 cultivated sunflower assembly was therefore used as a reference, and patterns of interactions were compared between samples. Raw sequence data was trimmed for enzyme cut site and base quality using the tool *trim* in the package HOMER[37] (v4.10) with the following flags ` -3 GATC -mis 0 -matchStart 20

-min 20 -q 15`. Trimmed data were then aligned to the HA412-HOv2 reference genome using NextGenMap[13] (v0.5.4) and interactions were quantified using the calls `makeTagDirectory -tbp 1 -mapq 10` and `analyzeHiC -res 1000000 -coverageNorm` from HOMER. This removes PCR duplicates based on mapping location, requires reads to have ≥10 mapping quality and normalizes interactions in 1 Mbp windows based on the total number of interactions. To determine which haploblocks differ between samples, aligned sequence data and samtools mpileup[15] were used to genotype diagnostic markers and call genotype for each haploblock, as described above.

Interpretation of the HiC patterns was sometimes complicated by the presence of putative structural differences between the genome of *H. petiolaris* and that of the HA412-HOv2 reference assembly against which reads from the HiC libraries were mapped. To determine what HiC patterns would be expected in those situations if haploblocks are associated to large inversions, we simulated an interaction matrix where interactions between windows linearly decayed based on distance. We then flipped window ordering within a region to simulate an inversion, and compare the interaction matrices with the original and flipped ordering. We used these basic HiC simulations to produce possible rearrangements between the haploblocks in *H. petiolaris* and the *H. annuus* reference that fit the observed HiC interaction patterns for three representative haploblocks (Extended Data Fig. 7b).

**Haploblock phenotype and environment associations**

In order to control for the effect of haploblocks on population structure, a variant file was created with all haploblock regions removed; both sites within haploblock regions and sites in close linkage (vcftools[25] v0.1.14, $R^2 > 0.5$) with haploblock genotypes were removed, in order to make sure that sites that were physically within the haploblock region were removed even if they were placed elsewhere due to reference differences. This haploblock-removed version of the genotype file was used for calculating PCA and kinship for EMMAX and the genetic covariance matrix for BayPass.

GWA analyses were performed using EMMAX[27] (v07Mar2010) for all traits measured in the common garden experiment (Supplementary Table 1). For all runs, the first three principal components (PCs) were included as covariates, as well as a kinship matrix calculated from the haploblock-removed genotype table. Environmental associations were run using BayPass[29] as previously described (see section "Genome-environment association analyses"), except that the 10,000 SNPs used to estimate population structure were drawn from the haploblock-removed dataset.

**Haploblocks phylogenies and dating**

A phylogenetic approach was used to determine the divergence time between haploblocks alleles. For each haploblock, five samples homozygous for each haploblock allele were chosen (defined as having >85% SNP ancestry from one haploblock allele). Two random samples from the other (sub)species, as well as two perennial samples (*H. grosseserratus* and *H. divaricatus*) were included in the analyses. For *H. petiolaris*, subsp. *petiolaris* and subsp. *fallax* were included in the same phylogeny if a haploblock was segregating in both. All genes within the haploblock in the HA412-HOv2 genome annotation were extracted, and the corresponding gene regions in the XRQv1 assembly were identified using a list of one-to-one orthologs between the two assemblies, created using Swiftortho[38]. For each gene, gVCF files were created from BAM files of the samples with GATK's (v4.0.6.0) HaplotypeCaller and gene sequences in FASTA format were generated using a custom Perl script. Haploblocks with more than 100 genes were down-sampled to 100 genes to reduce computing time.

### References

1 Schindelin, J. *et al.* Fiji: an open-source platform for biological-image analysis. *Nat. Methods* **9**, 676 (2012).

2 Schneider, C. A., Rasband, W. S. & Eliceiri, K. W. NIH Image to ImageJ: 25 years of image analysis. *Nat. Methods* **9**, 671 (2012).

3 Rodríguez, G. R. *et al.* Tomato Analyzer: a useful software application to collect accurate and detailed morphological and colorimetric data from two-dimensional objects. *JoVE*, e1856 (2010).

4 Murray, M. G. & Thompson, W. F. Rapid isolation of high molecular weight plant DNA. *Nucleic Acids Res.* **8**, 4321-4325 (1980).

5 Zeng, J., Zou, Y., Bai, J. & Zheng, H. Preparation of total DNA from recalcitrant plant taxa. *Acta Bot. Sin.* **44**, 694-697 (2002).

6 Rowan, B. A., Patel, V., Weigel, D. & Schneeberger, K. Rapid and inexpensive whole-genome genotyping-by-sequencing for crossover localization and fine-scale genetic mapping. *G3 (Bethesda)* **5**, 385-398 (2015).

7 Rohland, N. & Reich, D. Cost-effective, high-throughput DNA sequencing libraries for multiplexed target capture. *Genome Res.* **22**, 939-946 (2012).

8 Staton, S. E. *et al.* The sunflower (*Helianthus annuus* L.) genome reflects a recent history of biased accumulation of transposable elements. *Plant J.* **72**, 142-153 (2012).

9 Shagina, I. *et al.* Normalization of genomic DNA using duplex-specific nuclease. *Biotechniques* **48**, 455-459 (2010).

10 Matvienko, M. *et al.* Consequences of normalizing transcriptomic and genomic libraries of plant genomes using a duplex-specific nuclease and tetramethylammonium chloride. *PLoS One* **8**, e55913, doi:10.1371/journal.pone.0055913 (2013).

11 Bolger, A. M., Lohse, M. & Usadel, B. Trimmomatic: a flexible trimmer for Illumina sequence data. *Bioinformatics* **30**, 2114-2120 (2014).

12 Badouin, H. *et al.* The sunflower genome provides insights into oil metabolism, flowering and Asterid evolution. *Nature* **546**, 148-152 (2017).

13 Sedlazeck, F. J., Rescheneder, P. & von Haeseler, A. NextGenMap: fast and accurate read mapping in highly polymorphic genomes. *Bioinformatics* **29**, 2790-2791 (2013).

14 Li, H. A statistical framework for SNP calling, mutation discovery, association mapping and population genetical parameter estimation from sequencing data. *Bioinformatics* **27**, 2987-2993 (2011).

15 Li, H. *et al.* The Sequence Alignment/Map format and SAMtools. *Bioinformatics* **25**, 2078-2079 (2009).

16 Picard Tools, http://broadinstitute.github.io/picard/ (2019).

17 Tarasov, A., Vilella, A. J., Cuppen, E., Nijman, I. J. & Prins, P. Sambamba: fast processing of NGS alignment formats. *Bioinformatics* **31**, 2032-2034 (2015).

18 Poplin, R. *et al.* Scaling accurate genetic variant discovery to tens of thousands of samples. *BioRxiv*, doi:10.1101/201178 (2017).

19 Datta, K., Gururaj, K., Naik, M., Narvaez, P. & Rutar, M. GenomicsDB: Storing Genome Data as Sparse Columnar Arrays. *White Paper*, https://www.intel.com/content/dam/www/public/us/en/documents/white-papers/genomics-storing-genome-data-paper.pdf (2017).

20 Weng, M.-L. *et al.* Fine-grained analysis of spontaneous mutation spectrum and frequency in Arabidopsis thaliana. *Genetics* **211**, 703-714 (2019).

21 Zhang, W. *et al.* Comparing genetic variants detected in the 1000 genomes project with SNPs determined by the International HapMap Consortium. *Journal of genetics* **94**, 731-740 (2015).

22 Hubner, S. *et al.* Sunflower pan-genome analysis shows that hybridization altered gene content and disease resistance. *Nat. Plants* **5**, 54-62 (2019).

23 Köster, J. & Rahmann, S. Snakemake—a scalable bioinformatics workflow engine. *Bioinformatics* **28**, 2520-2522 (2012).

24 Li, H. Aligning sequence reads, clone sequences and assembly contigs with BWA-MEM. *ArXiv*, arXiv:1303.3997v1302 (2013).

25 Danecek, P. *et al.* The variant call format and VCFtools. *Bioinformatics* **27**, 2156-2158 (2011).

26 Browning, B. L., Zhou, Y. & Browning, S. R. A One-Penny Imputed Genome from Next-Generation Reference Panels. *Am. J. Hum. Genet.* **103**, 338-348 (2018).

27 Kang, H. M. *et al.* Variance component model to account for sample structure in genome-wide association studies. *Nat. Genet.* **42**, 348-354 (2010).

28 Grimm, D. G. *et al.* easyGWAS: A Cloud-Based Platform for Comparing the Results of Genome-Wide Association Studies. *Plant Cell* **29**, 5-19 (2017).

29 Gautier, M. Genome-Wide Scan for Adaptive Divergence and Association with Population-Specific Covariates. *Genetics* **201**, 1555-1579 (2015).

30 Jeffreys, H. Theory of Probability. Oxford University Press, London/New York/Oxford (1961).

31 Zheng, X. *et al.* A high-performance computing toolset for relatedness and principal component analysis of SNP data. *Bioinformatics* **28**, 3326-3328 (2012).

32 Hartigan, J. A. & Wong, M. A. Algorithm AS 136: A K-Means Clustering Algorithm. *J. R. Stat. Soc. C Appl. Stat.* **28**, 100-108 (1979).

33 R Core Team. R: A language and environment for statistical computing, https://www.R-project.org/ (2019).

34 Belton, J. M. *et al.* Hi-C: a comprehensive technique to capture the conformation of genomes. *Methods* **58**, 268-276 (2012).

35 Marie-Nelly, H. *et al.* High-quality genome (re)assembly using chromosomal contact data. *Nat. Commun.* **5**, 5695 (2014).

36 Sunflower Genome Database, https://www.sunflowergenome.org/ (2019).

37 Heinz, S. *et al.* Simple combinations of lineage-determining transcription factors prime cis-regulatory elements required for macrophage and B cell identities. *Mol. Cell* **38**, 576-589 (2010).

38 Hu, X. & Friedberg, I. SwiftOrtho: a Fast, Memory-Efficient, Multiple Genome Orthology Classifier. *BioRxiv*, 543223, doi:10.1101/543223 (2019).
